# The Wobble Uridine tRNA Writer MnmA Shapes Codon-Dependent Stress Response Systems

**DOI:** 10.64898/2026.08.15.745044

**Authors:** Humphrey C. Omeoga, Dylan Ehrbar, Chetna Mathur, Cady Roselli, Katherine Urner, Evan T. Davis, Rizwan Ahmad, Agnieszka Dziergowska, Qishan Lin, Peter C. Dedon, Jia Sheng, Thomas J. Begley

## Abstract

*Escherichia coli* uses wobble uridine (U34) modifications to tune codon decoding, but how individual tRNA writer enzymes shape gene expression remains unclear. Here, we identify MnmA, the U34 thiolation enzyme for tRNALys, tRNAGln, and tRNAGlu, as a central regulator linking codon-directed translation to regulon control and stress response. Loss of MnmA depleted s^2^U-dependent wobble modifications, preventing geranyl-(ges^2^U) and seleno-(se^2^U)-based modifications, causing growth defects, reduced catalase activity, and multi-level gene expression dysregulation. The Δ*mnmA* cells showed broad adaptive transcriptional reprogramming associated with RpoS- and OxyR-regulated pathways, which was accompanied by compromised protein output. Endogenous and tagged-protein analyses revealed specific impairment of transcriptional regulators, adaptive and detoxification proteins, including RpoS, OxyR, FliA, KatE, and KatG. Polysome profiling and polysome-associated RNA sequencing showed that MnmA deficiency globally reduces translational capacity and uncouples mRNA abundance from translational efficiency, which is exacerbated during oxidative stress. We developed genome-wide codon-usage mapping analytics to identify five codon-defined gene clusters, with specific clusters enriched for Lys, Gln, and Glu codons disproportionately affected by MnmA loss. Together, these findings support that wobble uridine thiolation and downstream modifications pair with corresponding codon architecture to coordinate the translation of regulon controllers and stress-response networks linked to bacterial fitness.

**Graphical Abstract:** 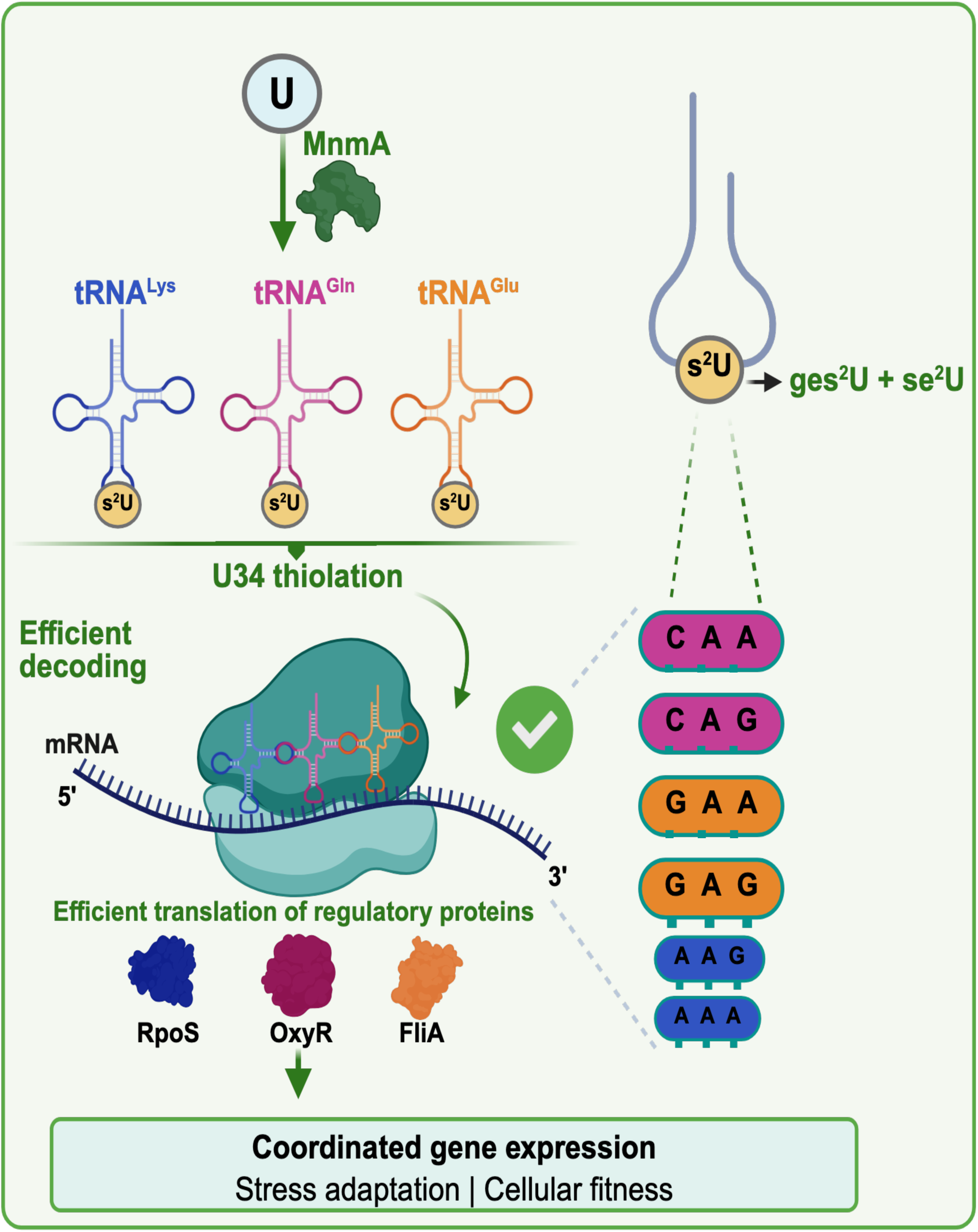

## Introduction

Transfer RNA (tRNA) contains over 170 posttranscriptional modifications, with many occurring at the wobble uridine (U34) position (1), a major hotspot of enzyme-catalyzed chemical diversity written and edited by tRNA modification systems (2,3). tRNA modifications regulate anticodon-codon interactions, translational speed, and fidelity by affecting ribosomal binding and frameshifting (4,5). tRNA modifications also contribute to stress responses through selective translation of Modification Tunable Transcripts (MoTTs), which have distinct codon usage biases (6–8). In Mycobacterium, the wobble modification (cmo^5^U) increases under hypoxia to regulate translation of the DosR/DosS mRNA (9). In *E. coli*, the tRNA 2’O-methylation (Gm18) has increased prevalence under stressed conditions (10), while in yeast, the wobble tRNA modifications 5-methoxycarbonylmethyluridine (mcm^5^U34) and 5-methylcytidine (m^5^C34) regulate the translation of DNA damage response and ribosomal genes with specific codon biases.

Wobble U34 modifications are strongly linked to oxidative stress response and the detoxification of reactive oxygen species (ROS) (11–13). ROS includes superoxide anions (O₂⁻), hydroxyl radicals (OH•), and hydrogen peroxide (H₂O₂), and all are by-products of aerobic metabolism (14). While basal ROS levels function as signaling molecules (14,15), excess ROS can disrupt cellular homeostasis and damage lipids, proteins, nucleic acids and iron–sulfur clusters (14,16,17), promoting mutagenesis and cell death (18–20). ROS can be counteracted by an integrated defense network comprising catalases, peroxidases, and superoxide dismutases (SODs) (14,17,21,22). Consistent with a functional connection between tRNA modifications and effective stress responses, disruption of the wobble U34-modifying enzyme GidA alters catalase activity in *Pseudomonas aeruginosa*, implicating wobble uridine pathways in bacterial ROS detoxification (23).

In *E. coli,* wobble U34 chemistry is coordinated by a set of enzymes that includes MnmA, MnmC (24–26), MnmE/G (27–31), and MnmH (32–34). The mnm-enzymes catalyze modifications including thiolation, methylaminomethylation, carboxymethylaminomethylation, geranylation, and selenation. MnmA catalyzes 2-thiolation (s^2^U) modifications at the wobble U34 (s^2^U) position of tRNA for lysine, glutamine, and glutamic acid. 2-thiolation (s^2^U) modification is crucial for *E. coli* cell growth, proliferation (35), and morphology (36,37). The MnmA-catalyzed s^2^U modification promotes tRNA interaction with A-ending codons (38) and serves as the substrate for the se^2^U- and ges^2^U-catalyzing enzyme MnmH (SelU). Loss of MnmA disrupts multiple wobble U34 modifications and has the potential to broadly impact translational regulation. Consistent with this role, MnmA deficiency reduces RpoS levels through impaired translation (38). RpoS is a global regulator of the stress response, controlling genes required for stationary-phase adaptation and oxidative stress resistance, including catalase KatE (39,40), and regulating over 150 genes in response to growth phase and environmental stress (21,41). Oxidative stress responses are further mediated by SoxRS and OxyR, which regulate expression of SODs (SodA, SodB*, and* SodC) (42,43), which convert superoxide into H₂O₂ for subsequent detoxification by catalases (KatG and KatE) (21,41,44–46). Together, these pathways maintain redox homeostasis under fluctuating conditions.

Here we investigate the role of MnmA in regulating codon-dependent translation, regulon controllers and stress-responsive gene expression. Using genetic, molecular, and systems-based approaches, we show that MnmA deficiency results in loss of s^2^U-dependent tRNA modifications, growth defects, increased sensitivity to oxidative stress, and a regulon-specific reprogramming of transcription. Notably, loss of MnmA impairs the translation of multiple transcription factors (RpoS, OxyR, and FliA), leading to dysregulation of stress-response and ROS detoxification pathways. Polysome-based measures revealed widespread dysregulation of protein synthesis in the absence of MnmA. We developed multi-signature codon cluster approaches to analyze the resulting translational efficiency data. We show that the *E. coli* genome has five distinct clusters of codon-defined genes and links MnmA to multiple codon-specific clusters. Together, our findings reveal MnmA as a central regulator of gene regulation with links to codon usage and provide a framework for understanding how wobble-uridine chemistry shapes bacterial physiology at the systems level.

## Materials and Methods

### Bacterial strains and growth conditions

*E. coli* wild-type K-12 BW25113 and the corresponding Δ*mnmA* mutant were obtained from the Keio Collection (47). Bacterial cells were transformed with plasmid pD444-SR (ATUM Biosciences, Newark, CA, USA) by the CaCl_2_ method and selected overnight at 37 °C on LB agar containing ampicillin (100 µg/mL). Transformants were grown overnight at 37 °C and 220 rpm in a shaker incubator (G-25, 281931, New Brunswick Scientific Co., Edison, NJ, USA), then diluted 1:1000 in LB and grown to OD_600_ values of 0.35–0.40 or 0.90–1.10, corresponding to exponential- and stationary-phase cultures, respectively.

### Catalase activity

Catalase activity was determined as described by Iwase et al. (48). Equal volumes of exponential- or stationary-phase cell cultures were mixed with equal volumes of Triton X-100 (Sigma-Aldrich, Burlington, MA, USA) and 35% hydrogen peroxide (H_2_O_2_; w/v) (Thermo Scientific, catalog number 202460010, Fisher Scientific Company, LLC, Pittsburgh, PA, USA). The height of catalase-generated bubbles was measured with a ruler and used as a measure of catalase activity. Measurements were performed using three biological replicates.

### Bacterial cell growth and sensitivity assay

For colony-forming unit (CFU) estimation, cultures were serially diluted 11-fold in 5-fold increments, spotted onto LB agar plates containing ampicillin (100 µg/mL), and either left untreated or supplemented with H_2_O_2_ (3.5 mM). The spotted plates were allowed to dry for 30 min at room temperature and then incubated at 37 °C for 16–18 h. Growth curves were initiated by diluting overnight cultures 1:1000 into fresh LB medium. OD_600_ readings were taken every 30 min using a microplate reader (Agilent BioTek, Santa Clara, CA) until OD_600_ reached 0.90–1.10.

### Motility assay

*E. coli* strains were grown to exponential phase, left untreated or treated with 5 mM H_2_O_2_, and then used for motility assays. Motility assays were performed as described by Galvanin et al.(10), with minor modifications. Overnight cultures were diluted and grown to exponential phase, and 3 µL of exponential-phase culture was inoculated into semisolid LB agar (0.35%; wt/vol). Images were captured after 24 h of incubation at 37 °C, and the area of bacterial spread was measured.

### mRNA-sequencing

Exponential- or stationary-phase cultures were either left untreated or treated with 5 mM H_2_O_2_ for 30 min. Cells were harvested by centrifugation at 9,800 × g for 5 min in 15-mL centrifuge tubes containing a stop solution composed of 5% water-saturated phenol in ethanol (pH < 7.0). Total RNA was isolated using the hot-phenol method (49). Total RNA samples were quantified using a Qubit 4.0 Fluorometer (Life Technologies, Carlsbad, CA, USA), and RNA integrity was assessed with a 4200 TapeStation (Agilent Technologies, Palo Alto, CA, USA).

### Library Preparation and Sequencing

Samples with >10% DNA contamination were treated with TURBO DNase (Thermo Fisher Scientific, Waltham, MA, USA). RNA normalization was performed by adding the ERCC RNA Spike-In Mix (Cat. #4456740; Thermo Fisher Scientific) according to the manufacturer’s protocol. After DNase treatment, dual rRNA depletion was performed using the QIAGEN FastSelect rRNA HMR Kit (Qiagen, Germantown, MD, USA) according to the manufacturer’s protocol. RNA sequencing libraries were constructed using the NEBNext Ultra II RNA Library Preparation Kit for Illumina, following the manufacturer’s recommendations. In brief, enriched RNAs were fragmented for 15 min at 94 °C. Fragmented RNA was converted into complementary DNA (cDNA), and unique molecular identifiers (UMIs) were incorporated during reverse transcription. cDNA fragments were end-repaired and adenylated at their 3′ ends, and universal adapters were ligated to cDNA fragments, followed by index addition and library enrichment with limited-cycle PCR. The sequencing libraries were validated using the Agilent TapeStation 4200 (Agilent Technologies, Palo Alto, CA, USA) and quantified using the Qubit 4.0 Fluorometer (Thermo Fisher Scientific, Waltham, MA, USA) and quantitative PCR (KAPA Biosystems, Wilmington, MA, USA). Sequencing libraries were multiplexed, clustered on a flow cell, and sequenced on an Illumina NovaSeq 6000 instrument according to the manufacturer’s instructions. Samples were sequenced using a 2 × 150 bp paired-end configuration. Raw sequence data (.bcl files) generated by the sequencer were converted into FASTQ files and demultiplexed using Illumina’s bcl2fastq 2.17 software. One mismatch was allowed for index sequence identification. FASTQ files were aligned using STAR, and gene-level count files were generated. FASTQ files have been deposited in the NCBI database (GEO accession number: GSE330173), and differential gene expression was analyzed using Bash and RStudio (2023.06.2+561; Posit Software, PBC, Boston, MA). Volcano plots were generated using a fold-change cutoff of 1.50 and a p-value cutoff of 0.05. Gene Ontology (GO) analysis of upregulated and downregulated genes was performed using the clusterProfiler package (50).

### Protein quantification assay

Native RpoS and GAPDH were detected using primary antibodies from BioLegend (catalog number 663703, San Diego, CA, USA) and Invitrogen (catalog number MA5-15738, USA), respectively. *E. coli* cells were harvested by centrifugation and washed with 1× PBS to remove residual medium. Cells were lysed using lysis buffer (Thermo Scientific B-PER reagent, product number 78243; protease inhibitor, product number A32955; and DNase I from Fisher Scientific LLC, Pittsburgh, PA, USA) and incubated at room temperature for 30–60 min. Lysates were then centrifuged at 13,000 × g and 4 °C for 5 min, and the supernatant containing the protein lysate was collected and stored at −20 °C. Protein quantification was performed using the Pierce^TM^ BCA Protein Assay (product number 23227, Thermo Fisher Scientific, USA) according to the manufacturer’s protocol. Protein lysates were adjusted to 2.0 µg/µL, and 6 µg total protein was analyzed by WES (ProteinSimple, San Jose, CA, USA) using the manufacturer’s analysis plates (catalog no. SM-W004, ProteinSimple, San Jose, CA, USA).

### Polysome profiling

Polysome-associated mRNA was analyzed as previously described (51). In brief, *E. coli* WT and Δ*mnmA* cells were grown to exponential and stationary phases and either treated with H_2_O_2_ or left untreated. Cultures were transferred into 50-mL centrifuge tubes packed loosely with crushed ice. Tubes were quickly capped and centrifuged at 10,000 rpm for 5 min at 4 °C. Cell pellets were resuspended in chilled lysis buffer (10 mM Tris-HCl, pH 8.0, 10 mM MgCl_2_, 1 mg/mL lysozyme) and transferred to 1.5-mL Eppendorf tubes. Resuspended cells were flash-frozen in liquid nitrogen and thawed in a cool water bath with occasional tube inversion. The thawed cells were flash-frozen again, thawed, and 15 µL of 10% sodium deoxycholate was added to complete lysis. Lysed cells were centrifuged at 10,000 rpm for 10 min at 4 °C. The supernatant was transferred into a fresh 1.5-mL Eppendorf tube and stored on ice. Sucrose gradients (10%– 40%) containing 20 mM Tris-HCl, pH 7.8, 10 mM MgCl_2_, 100 mM NH_4_Cl, and 2 mM dithiothreitol (DTT) were prepared and stored at 4 °C. Equal volumes of cell supernatant were layered onto sucrose gradients, balanced to 0.001 g, and loaded into SW41 rotor buckets. Gradients were ultracentrifuged at 35,000 rpm for 3 h at 4 °C. With 50% sucrose as the push liquid, gradient fractionation was performed using the Teledyne ISCO Foxy JR fractionator as described by the manufacturer (Teledyne ISCO, Lincoln, NE, USA). Mean values from three biological replicates were calculated, and the area under the curve (AUC) for polysome:70S and disome ratios was calculated using the trapezoidal rule. Statistical significance was assessed using Student’s *t*-test in GraphPad Prism v10 (La Jolla, CA, USA).

### Polysome RNA isolation and NGS RNA-seq

Total RNA was extracted separately from individual polysome fractions using a phenol-chloroform RNA extraction protocol adapted from Rhodius and Wade (49). Library preparation and STAR alignment were performed as described above, and the data have been deposited in the NCBI Gene Expression Omnibus (GEO; accession number GSE330174).

### His-tagged gene design and WES-based detection

To detect proteins of interest, the coding sequences of *rpoS*, *oxyR*, *katE*, *katG*, *ahpC*, and *fliA* were engineered to encode C-terminal 6×His tags and cloned into the pD444-SR plasmid purchased from ATUM. The plasmids were then transformed into wild-type and mutant *E. coli* cells and selected on ampicillin-containing medium. Single colonies of transformed cells were inoculated and grown overnight. Cultures were then diluted and grown to the exponential phase. Exponential-phase cells (OD_600_ of ∼0.4) were either left untreated or treated with 5 mM H_2_O_2_ for 30 min, then harvested for total protein extraction and quantification. After protein extraction and quantification, protein detection was performed using a 1:10,000 anti-His-tag horseradish peroxidase-conjugated antibody (MAB050H, R&D Systems) on the WES ProteinSimple system.

### Quantification of tRNA modifications

tRNA modifications were quantified by liquid chromatography-coupled tandem mass spectrometry (LC-MS/MS) as described by Chan et al. (52). Small RNAs were isolated from Δ*mnmA* and wild-type cells at different growth phases and under different treatment conditions using the Thermo Fisher PureLink^TM^ miRNA Isolation Kit (cat. #K157001) according to the manufacturer’s protocol. Purified small RNA samples were digested at room temperature using the Nucleoside Digestion Mix (Catalog No. M0649S, New England BioLabs, Beverly, MA, USA) and stored at 4 °C. Following small-RNA digestion, modified nucleosides were analyzed on a Waters XEVO TQ-S^TM^ electrospray triple-quadrupole mass spectrometer (Waters, Milford, MA, USA) coupled to an ACQUITY I-Class ultra-performance liquid chromatography system, as described by Basanta-Sanchez et al. (53). The capillary voltage was set to 1.0 kV, with an extraction cone voltage of 14 V. Nitrogen flow was maintained at 1000 L/h, and the desolvation temperature was 500 °C. The cone gas flow was set to 150 L/h and the nebulizer pressure to 7 bars. Each nucleoside modification was characterized by single infusion in positive-ion mode over an *m/z* range of 100–500 amu. Further nucleoside characterization was performed using Waters software as part of the IntelliStart MS/MS method development program, in which a ramp of collision and cone voltages was applied to determine optimal collision-energy parameters for daughter ions. Detected modifications were identified and quantified by comparison with reference standards. Stable isotope-labeled guanosine [^13^C][^15^N]-G (1 pg/µL) (cat. #CNLM-3808-CA-5, Cambridge Isotope Laboratories, Inc., Tewksbury, MA, USA) was used as an internal standard. Chemically synthesized mnm^5^U and mnm^5^s^2^U were used as reference standards to validate and quantify modification levels detected in experimental samples.

### Codon usage analysis and Gaussian mixture modeling

Total codon frequencies were calculated for all annotated protein-coding genes in the *Escherichia coli* genome by counting the 61 codons per coding sequence and converting the counts to within-gene relative frequencies. Total codon frequencies were Z-score normalized across the genome to mitigate gene-length effects and enable direct comparisons among genes. To identify genome-wide patterns of codon usage, Gaussian mixture models (GMMs) were fit to the normalized codon-frequency matrix using an expectation-maximization framework, testing a range of component numbers and covariance structures (54). Model selection was performed using the Bayesian Information Criterion (BIC) (55,56), which balances goodness of fit and model complexity. The optimal model was used to define discrete codon-usage clusters across the genome. The top-ranked model for total codon frequency was the VVE model with five clusters (VVE5).

### Translational efficiency mapping

Translational efficiency (TE) was estimated using a modified version of the method described by Cui et al. (57) as the ratio of polysome-associated mRNA abundance to total mRNA abundance. Differential expression values from polysome and crude lysate RNA-seq comparisons were merged by gene name, and genes were classified according to TE direction and magnitude using predefined log₂ fold-change and significance thresholds (|log2FC| ≥ 1.50 and padj ≤ 0.05). For quadrant-based analyses, genes were assigned to one of four categories based on TE and lysate log₂ fold change: TE_up_mRNA_down, TE_up_mRNA_up, TE_down_mRNA_down, and TE_down_mRNA_up. For collapsed translational classes, TE-up genes were defined as the union of TE_up_mRNA_down and TE_up_mRNA_up, and TE-down genes as the union of TE_down_mRNA_down and TE_down_mRNA_up.

To determine whether specific codon-usage clusters were overrepresented within TE-defined gene groups, Fisher’s exact test was applied separately to each cluster, using 2 × 2 contingency tables that compared the number of genes from each translational class within a cluster with the number expected based on the genome-wide cluster distribution. Odds ratios (ORs) and raw P-values were calculated for each comparison, and multiple-testing correction was performed using the Benjamini–Hochberg false discovery rate procedure.

Codon signature analyses were performed by mapping TE-defined genes onto the genome-wide codon usage framework generated from the Z-score-normalized codon matrix. Codon signature values were therefore derived from the same globally normalized codon space used for Gaussian mixture modeling. For each TE-defined group, mean codon Z-scores were computed across all codons using the globally normalized codon matrix. From this common codon space, summary metrics were derived by grouping codons based on third-position nucleotide identity (A/U-ending versus G/C-ending) or by examining specific wobble uridine–sensitive synonymous codons (AAA/AAG, GAA/GAG, and CAA/CAG). Group-level codon signatures were summarized as the mean ± 95% confidence interval, with confidence intervals estimated by bootstrap resampling of genes within each class.

## Results

### MnmA writer-deficient cells exhibit impaired growth, reduced viability, and compromised catalase activity

We assessed the physiological consequences of disrupting Mnm enzymes by initially comparing the exponential-phase growth phenotypes of individual mutants (Δ*mnmA*, Δ*mnmC*, Δ*mnmE*, Δ*mnmG*, and Δ*mnmH*). Among these strains, Δ*mnmA* cells displayed the most severe growth defect, whereas Δ*mnmE* and Δ*mnmG* cells showed only mild growth impairment **(Figure 1A)**. Consistent with this growth phenotype, Δ*mnmA* cells also exhibited a significant reduction in viability upon entry into stationary phase, as measured by CFU assays **(Figure 1B)**. Complementation of the Δ*mnmA* strain with a plasmid expressing *E. coli mnmA* restored growth to near wild-type levels in rich medium **(Figure 1C)**.

**Figure 1.**
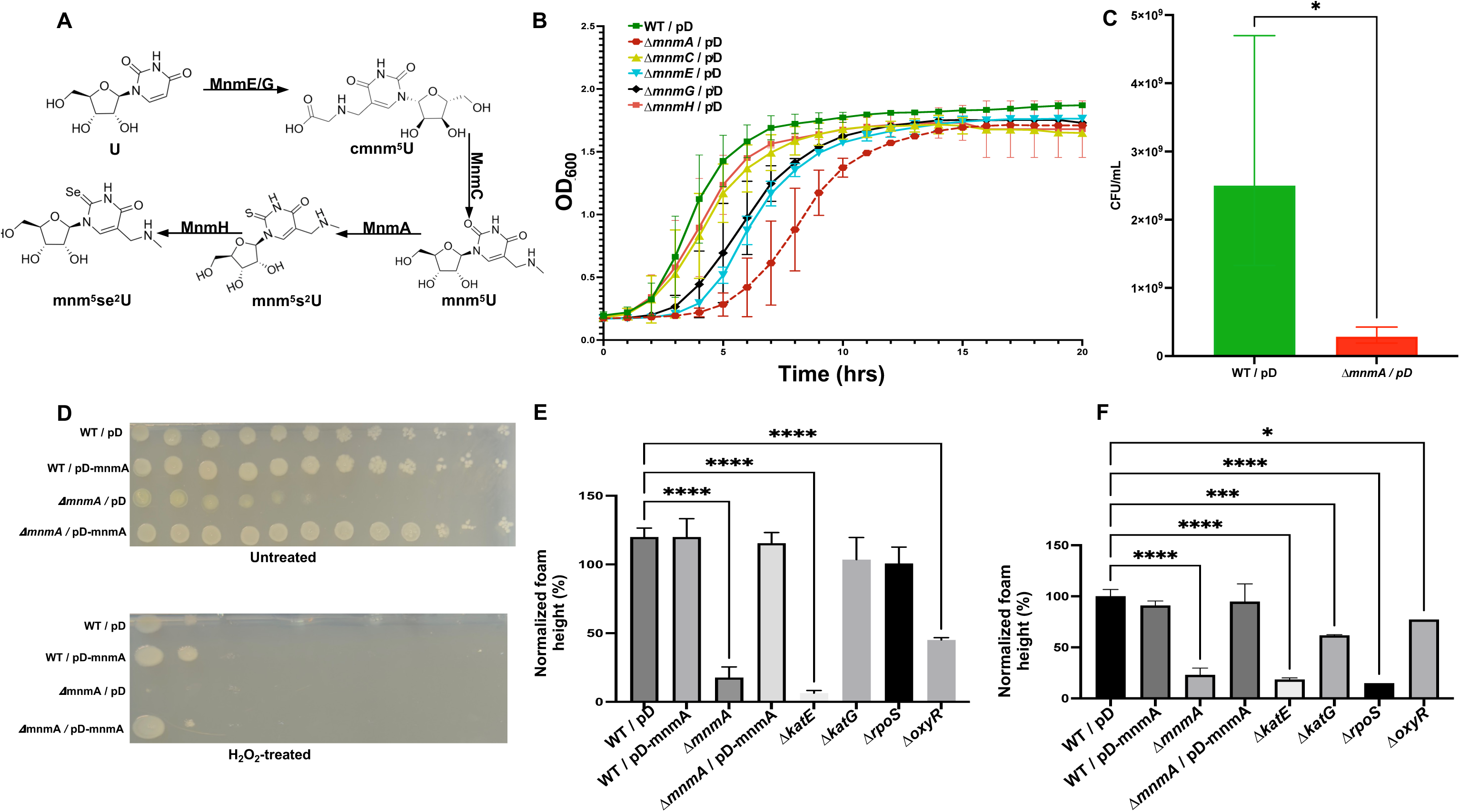
The MnmA promotes cell growth and catalase activity. **(A)** Schematic of the wobble uridine (U34) modification pathway showing the chemical structures of U34 modifications generated by MnmE-MnmG, MnmC, MnmA and MnmH, in tRNAs for lysine, glutamine and glutamate. **(B)** Growth curves of WT and the indicated mnm-deletion mutants (Δ*mnmA,* Δ*mnmC,* Δ*mnmE,* Δ*mnmG,* and Δ*mnmH*; n = 3 biological replicates). **(C)** Colony-forming unit (CFU) analysis of WT and Δ*mnmA* cells during stationary phase (n = 3 biological replicates). **(D)** A sensitivity assay was performed on WT, WT/pD-mnmA, Δ*mnmA,* and Δ*mnmA*/pD-mnmA, either left untreated or treated with 5 mM H_2_O_2_. **(E, F)** Catalase activity was measured in WT, WT/pD-mnmA, Δ*mnmA,* Δ*mnmA*/pD-mnmA, and the control strains Δ*katE,* Δ*katG,* Δ*oxyR,* and Δ*rpoS* during **(E)** exponential phase and **(F)** stationary phase. Statistical significance was determined from three biological replicates (n = 3) by one-way ANOVA (* p < 0.05, ** p < 0.01, **** p < 0.0001).

Because tRNA modification mutants commonly display stress-associated phenotypes (6,8,23,58), we next assessed sensitivity to H_2_O_2_ in Δ*mnmA* cells, which were the most stressed among the strains based on growth phenotypes. MnmA-deficient cells showed modest sensitivity to oxidative stress relative to wild-type cells, and this phenotype was rescued by *mnmA* complementation **(Figure 1D)**. We also extended this analysis to other members of the mnm-enzyme cluster **(Supplementary Figures S1A-B)**. Because catalase activity is central to ROS detoxification, it was measured in wild-type, Δ*mnmA*, and control strains during exponential and stationary phases. Δ*mnmA* cells showed a significant reduction in catalase activity relative to wild-type cells in both growth phases **(Figures 1E, F)**, and complementation restored catalase activity to wild-type levels. The expected reductions in catalase activity in Δ*katE*, Δ*katG*, Δ*oxyR,* and Δ*rpoS* strains placed the Δ*mnmA* phenotype within established oxidative stress response pathways **(Figure 1F)**. To confirm the tRNA modification defect, we show that Δ*mnmA* cells have ∼10-fold higher levels of mnm^5^U and ∼100-fold lower levels of mnm^5^s^2^U, with the latter essentially zero at the detection limits. In addition, we demonstrate that the s^2^U-based modification increases during the stationary phase compared with the exponential phase **(Supplementary Figures S2A-B)**. Together, these results identify MnmA-dependent wobble U thiolation as an important determinant of bacterial growth, stationary-phase survival, and oxidative stress defense.

### Loss of MnmA drives basal transcriptional reprogramming during exponential growth

To determine whether the physiological defects observed in MnmA-deficient cells were accompanied by transcriptional changes, we performed mRNA-seq on wild-type and Δ*mnmA* cells grown to exponential phase and exposed to H_2_O_2_. We compared untreated Δ*mnmA* cells with untreated wild-type cells to reveal extensive basal transcriptional dysregulation, with 194 genes downregulated and 438 upregulated, using an absolute log_2_ fold-change cutoff >1.50 and adjusted *p*-value ≤0.05 **(Figure 2A)**. Thus, MnmA deficiency is associated with broad transcriptional reprogramming even in the absence of external stress. Gene ontology analysis of downregulated transcripts showed significant enrichment for ribosomal genes, rRNA-associated processes, and translation-related pathways, whereas upregulated transcripts were enriched for glutathione metabolism, ABC transporters, and general metabolic functions **(Supplementary Figure S3)**.

**Figure 2.**
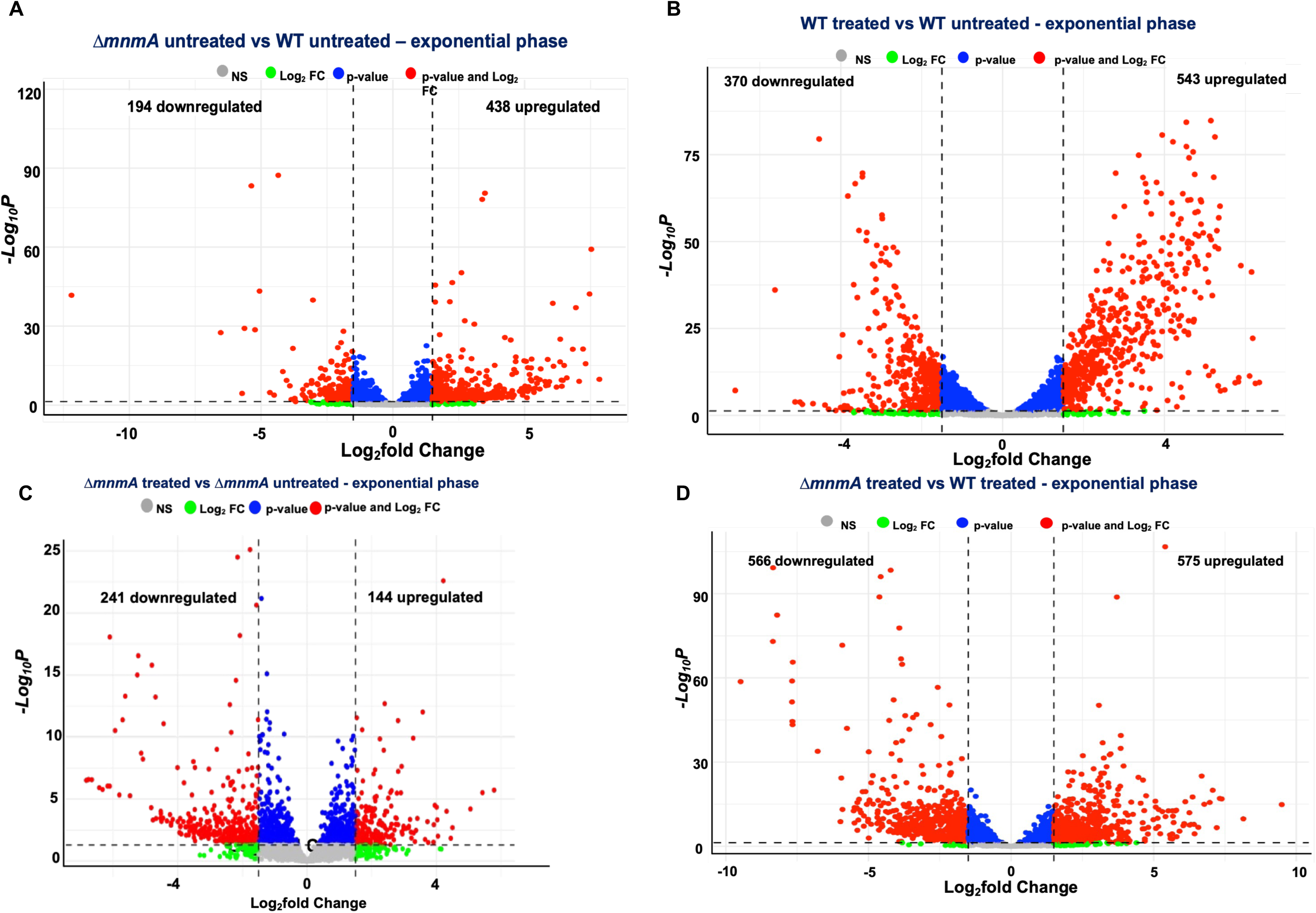
Basal and stress-induced transcription are globally altered in MnmA-deficient cells during the exponential phase. RNA from Δ*mnmA* untreated vs WT untreated cells in exponential and stationary phases was purified and analyzed by mRNA-seq. Enhanced volcano plot of the mRNA-seq data was plotted for **(A)** *ΔmnmA* untreated vs WT untreated – exponential phase, **(B)** WT treated vs WT untreated - exponential phase**, (C)** *ΔmnmA* treated vs *ΔmnmA* untreated - exponential phase, and **(D)** *ΔmnmA* treated vs WT treated - exponential phase.

### MnmA deficiency alters the transcription of oxidative stress systems in response to H_2_O_2_

We next examined how loss of MnmA altered the transcriptional response to oxidative stress. H_2_O_2_ exposure induced extensive transcriptional reprogramming in both wild-type and MnmA-deficient cells during exponential growth **(Figures 2B, C)**. However, direct comparison of H_2_O_2_-treated Δ*mnmA* cells with treated wild-type cells revealed substantial genotype-dependent differences, with 566 transcripts downregulated and 575 transcripts upregulated in the mutant background **(Figure 2D)**. Gene ontology analysis across strains and conditions further showed that, under basal conditions, motility-associated genes were enriched among transcripts upregulated in Δ*mnmA* cells **(Supplementary Figure S3)**, a pattern not observed in response to H_2_O_2_ for either genotype (**Supplementary Figures S4–S5).** A comparison of Δ*mnmA*-treated cells with wild-type-treated cells showed that, in the mutant, oxidative stress promoted upregulation of transcripts involved in ribosome assembly, translation, and organelle assembly **(Supplementary Figure S6)**. Heatmap visualization confirmed that untreated Δ*mnmA* cells exhibit a transcriptional profile distinct from that of untreated wild-type cells **(Figure 3A)**. Because motility-associated transcripts were elevated in Δ*mnmA* cells, we performed a swarming assay to test whether this transcriptional signature translated into increased motility. Despite the increased expression of motility genes, MnmA-deficient cells showed no significant increase in swarming but showed a significant decrease in response to oxidative stress **(Supplementary Figure S8)**. Together, these findings support that MnmA-deficient cells undergo extensive transcriptional remodeling during oxidative stress, but this response is qualitatively distinct from the wild-type program.

**Figure 3:**
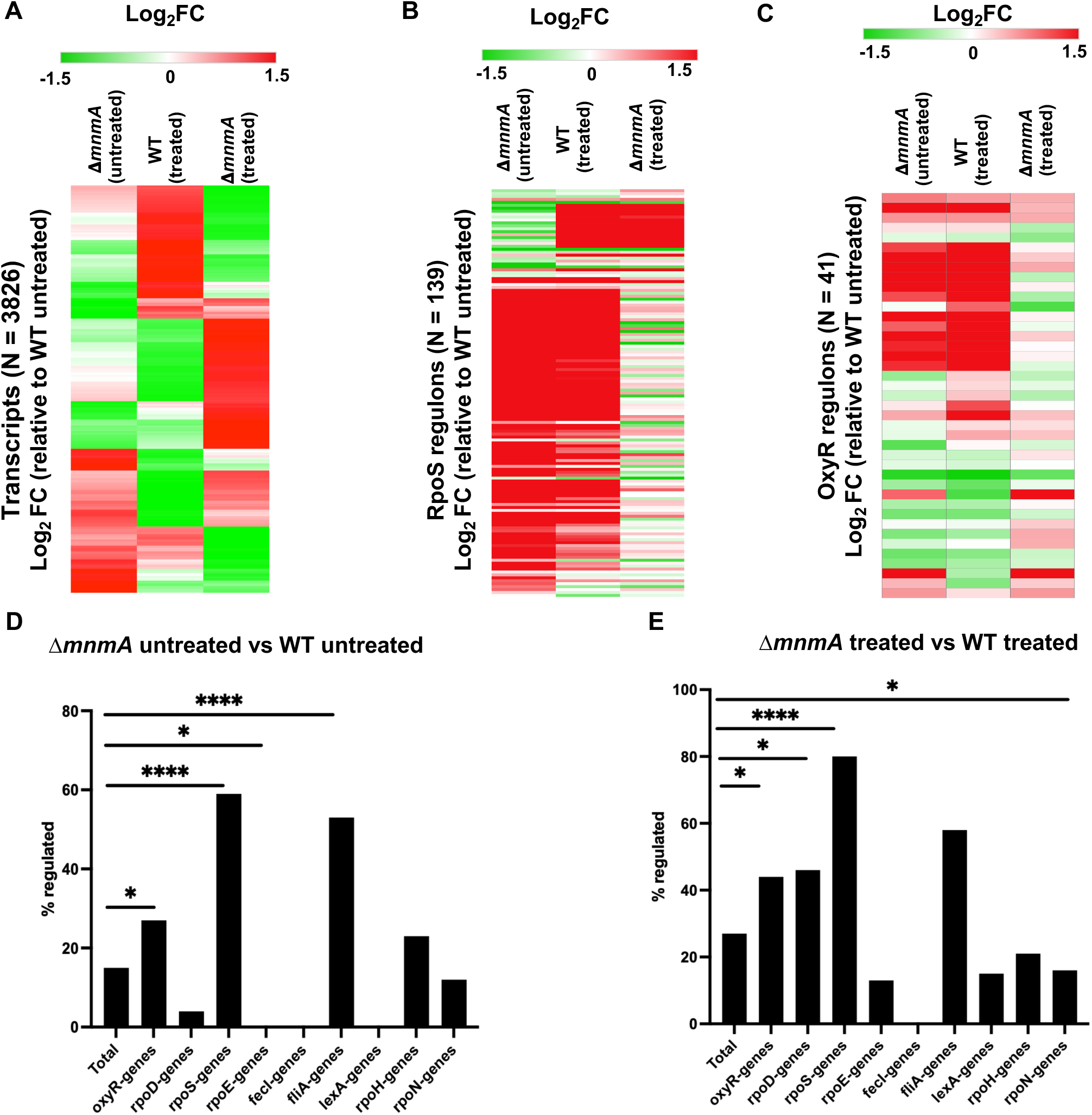
Heatmap visualizations of clustered genes and bar graphs of dysregulated regulons during the exponential growth phase. Differentially expressed genes are plotted across different cell conditions and compared to the WT-untreated condition. Values are represented (log_2_FC ≥ 1.50 and log_2_FC ≤ 1.50, *P_adj._* ≤ 0.05) for **(A)** all DEGs (N = 3,826), **(B)** RpoS-regulated (N = 139), and **(C)** OxyR-regulated transcripts (N = 41). Fisher’s exact test was used to assess significant dysregulation of the *E. coli* regulons, and the results were presented as bar graphs in **(D)** Δ*mnmA* untreated vs. WT untreated and **(E)** Δ*mnmA* treated vs. WT treated. (* *p < 0.05*, **** *p < 0.0001*).

### Stress-response regulons are transcriptionally dysregulated in Δ*mnmA* cells

We further investigated the transcriptional consequences of MnmA deficiency by assessing whether specific regulons were affected. We examined genes regulated by the major stress-associated transcription factors RpoS and OxyR. RpoS-regulated genes were upregulated in untreated MnmA-deficient cells (N = 78), resembling the transcriptional response seen in H_2_O_2_-treated wild-type cells **(Figure 3B)**. However, after oxidative stress, Δ*mnmA* cells induced only a small fraction of RpoS-dependent genes (N = 16), suggesting that MnmA-deficient cells enter a partially stress-activated state but have a reduced capacity to further remodel the RpoS regulon upon challenge. A similar pattern was observed for OxyR-regulated genes, including *katE, katG, ahpC, dps,* and components of the *suf* operon, which were elevated in untreated Δ*mnmA* cells and mirrored a stress-induced profile in wild-type cells **(Figure 3C)**. Beyond the RpoS- and OxyR-regulons, statistical analysis identified significant dysregulation of other regulon-associated genes across basal and oxidative stress conditions in MnmA-deficient cells **(Figures 3D, E; Tables 1 and 2; Supplementary Figure S7)**. In contrast, the stationary-phase transcriptomes showed fewer genotype-dependent differences, suggesting a more prominent role for MnmA during exponential growth **(Supplementary Figures S9–S13)**.

**Table 1:**
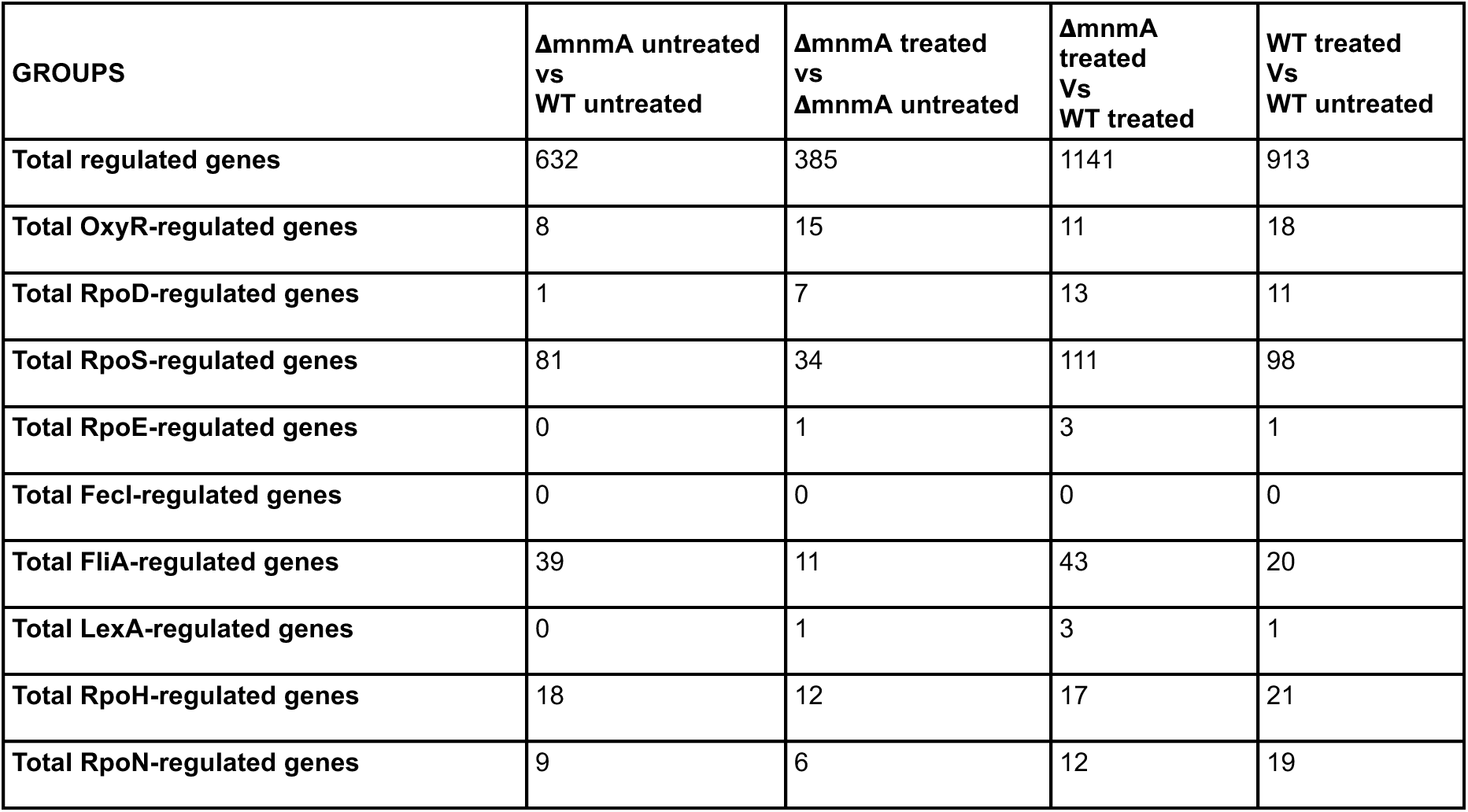
Dysregulated genes for some E. coli regulons in different groups.

**Table 2:**
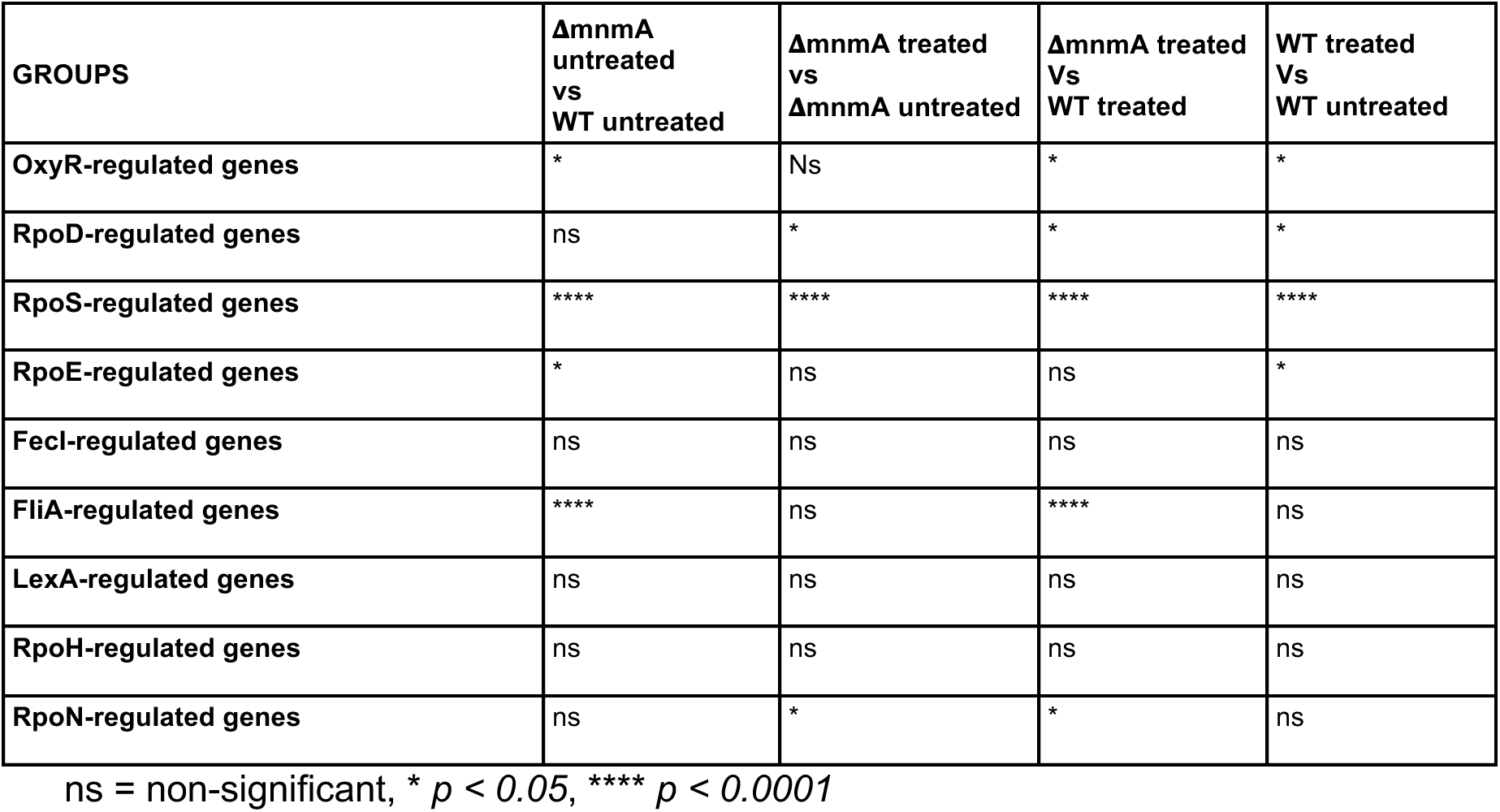
Statistical analysis of total dysregulated regulons for different groups.

### Endogenous RpoS protein levels are impaired in MnmA-deficient cells

Given the dysregulation of stress-response regulons, we next examined the expression of the master stress regulator, RpoS, in both the exponential and stationary phases, and of other *E. coli* stress-response proteins only in the exponential phase. In wild-type cells, RpoS protein levels increased in response to H_2_O_2_ in both exponential and stationary phases **(Figures 4A** and **4B)**. In contrast, MnmA-deficient cells showed little to no detectable RpoS protein under either basal or oxidative stress **(Figures 4A, B)** during the different growth phases. Because RpoS is normally induced under oxidative stress and in the stationary phase, the failure of MnmA-deficient cells to accumulate RpoS suggests a defect beyond environmental changes. These results are consistent with post-transcriptional regulation (59,60), and link MnmA-catalyzed s^2^U modification to RpoS protein expression as part of a broader disruption of bacterial gene expression control.

**Figure 4.**
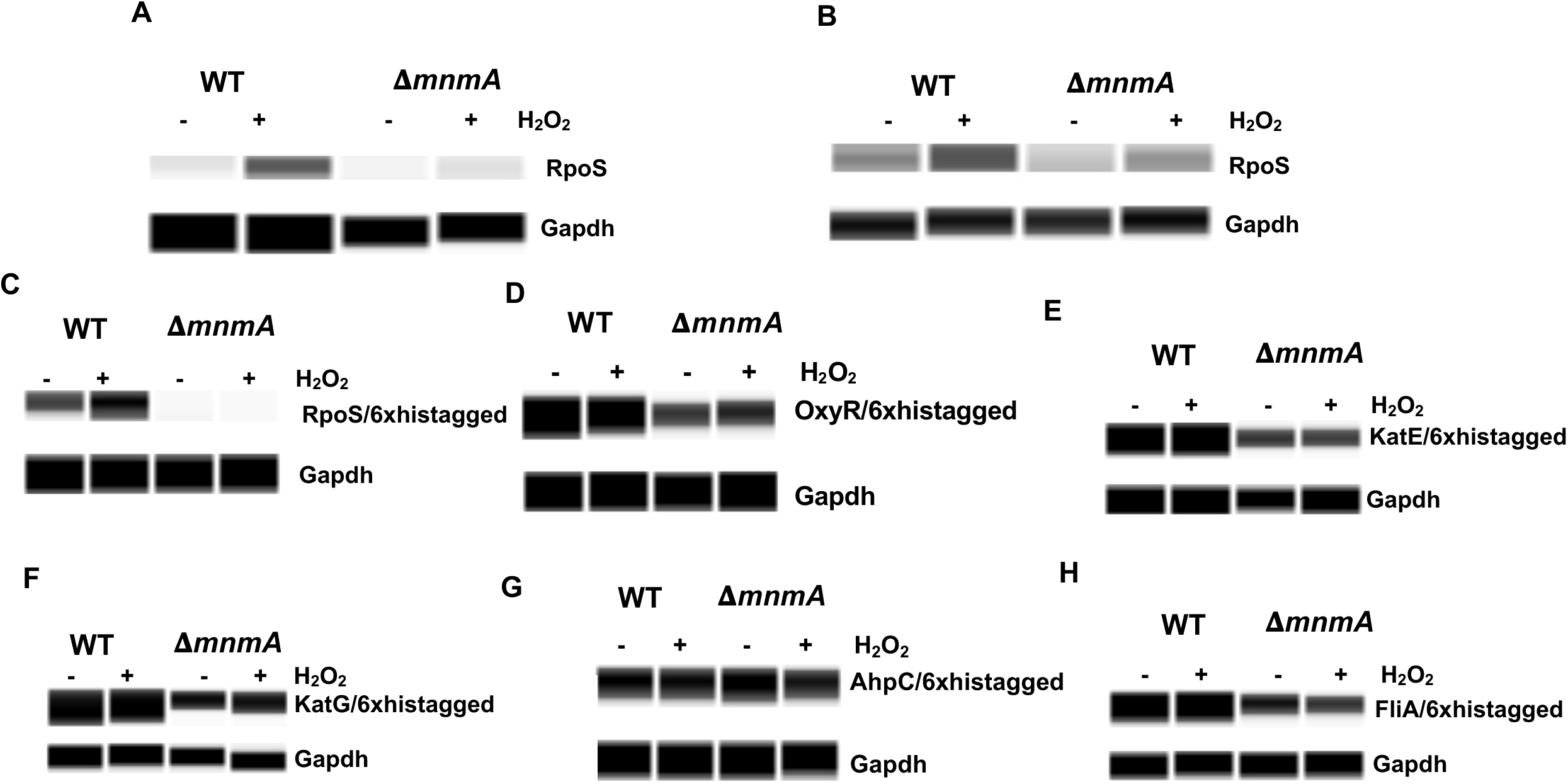
MnmA deficiency impairs the accumulation of endogenous RpoS and selected tagged stress-response proteins. **(A–B)** Endogenous RpoS levels in wild-type (WT) and Δ*mnmA* cells during exponential **(A)** and stationary **(B)** phases under untreated conditions or with 5 mM H₂O₂ for 30 min. **(C–H)** Protein levels for C-terminal 6xHis-tagged RpoS, OxyR, KatE, KatG, AhpC, and FliA in WT and Δ*mnmA* cells during the exponential phase under untreated conditions or after treatment with 5 mM H₂O₂. Protein abundance was measured using the ProteinSimple WES system.

### Impairment of specific stress-response protein levels in *mnmA* cells

We observed that endogenous RpoS protein levels were reduced in Δ*mnmA* cells, prompting us to investigate the expression of the C-terminal His-tagged transcription factors and oxidative stress detoxification enzymes: OxyR, RpoS, KatE, KatG, AhpC, GapA, and FliA. Tagged constructs were used because suitable antibodies were not available to the native proteins. FliA was included because our transcriptomic data showed that flagellar genes were dysregulated in the Δ*mnmA* cells **(Supplementary Table 1),** and our swarm assay shows a significant decrease in swarm activity in the MnmA-deficient cells when exposed to oxidative stress relative to the basal condition **(Supplementary Figures S8B-C).** RpoS, OxyR, KatE, KatG, and FliA protein levels were consistently reduced in MnmA-deficient cells relative to wild-type, while AhpC showed comparable expression in both genetic backgrounds **(Figures 4C-H)**. Our findings suggest that loss of MnmA impairs the production of many proteins with a subset of proteins involved in regulon control and stress response more severely affected.

### MnmA deficiency reduces polysome abundance and alters translational efficiency

Given the combined transcriptional dysregulation and impaired protein abundance in Δ*mnmA* cells, we next asked whether the MnmA deficiency broadly affects translation. To test this, we performed polysome profiling on wild-type and Δ*mnmA* cells under untreated and H_2_O_2_-treated conditions during exponential and stationary phases. Under basal exponential-phase conditions, Δ*mnmA* cells showed reduced polysome abundance relative to wild-type cells **(Figure 5A),** indicating an impaired capacity for translation. Polysome-associated RNA-seq further revealed widespread changes in ribosome-associated transcripts in *ΔmnmA* cells **(Figure 5B)**. Functional enrichment analysis showed that polysome-associated transcripts for iron-sulfur cluster binding, fermentation, and response to acidic stress were decreased in *ΔmnmA* cells, whereas those for cell motility, ribosome assembly, and organelle organization were increased **(Figure 5C)**. Translation efficiency (TE) analysis confirmed broad shifts in mRNA translation in *ΔmnmA* cells **(Figure 5D)**. Oxidative stress further reshaped the translational profile of MnmA-deficient cells, producing patterns distinct from the basal condition **(Supplementary Figures S14–S17).** H_2_O_2_ exposure caused a pronounced reduction in polysome abundance in the Δ*mnmA* cells relative to both untreated and treated cells **(Supplementary Figures S16A, B).** Under oxidative stress, TE analysis showed reduced translation of stress-response genes and increased translation of mRNAs associated with flagellar processes, chemotaxis, and motility in *ΔmnmA* cells compared with wild-type cells **(Supplementary Figure 17B)**. Stationary-phase RNA-seq showed fewer genotype-dependent transcriptional differences **(Supplementary Figures S9–S13)**, consistent with the more limited polysome-associated changes observed during stationary phase **(Supplementary Figures S18–S22).** Together, these results support the idea that MnmA deficiency reduces translational capacity and reshapes translational efficiency under both basal and oxidative-stress conditions.

**Figure 5.**
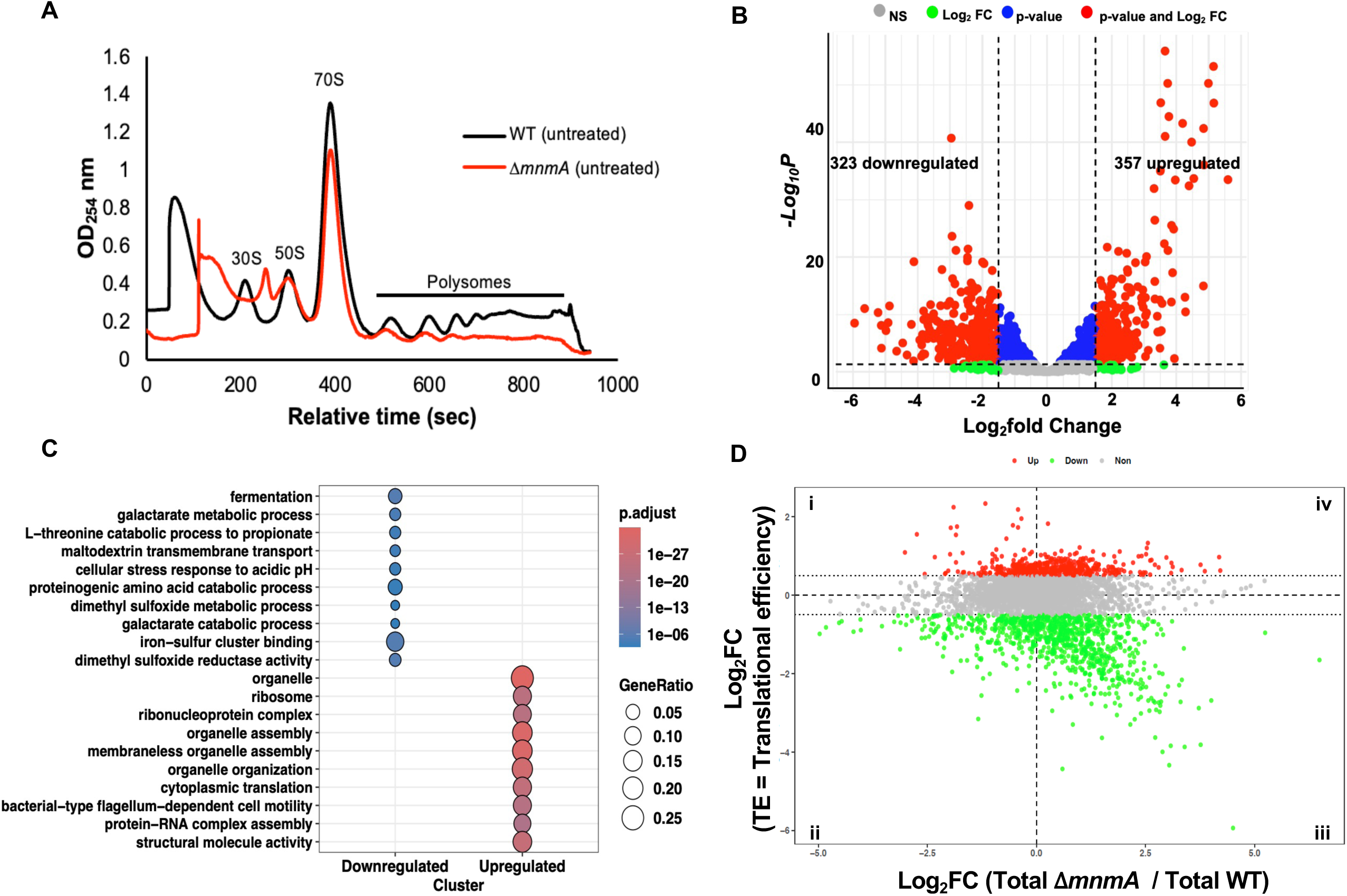
MnmA deficiency reduces polysome abundance and reshapes translational efficiency during exponential growth. **(A)** Polysome profiles of untreated exponentially growing WT and Δ*mnmA* cells. Absorbance at 254 nm (OD₂₅₄) is plotted across gradient fractions, with peaks corresponding to 30S, 50S, 70S, and polysome fractions. **(B)** Volcano plot showing differential abundance of polysome-associated transcripts in Δ*mnmA* relative to WT under untreated conditions. Significantly increased (357) and decreased (323) polysome-associated transcripts are indicated based on defined fold-change and adjusted *P*-value thresholds. **(C)** Gene Ontology enrichment analysis of differentially abundant polysome-associated transcripts using clusterProfiler. Dot size represents the gene ratio, and color indicates the adjusted *P*-value. **(D)** Translational efficiency (TE) analysis comparing TE versus total mRNA changes in Δ*mnmA* relative to WT cells. Genes are classified as TE-up (red), TE-down (green), or unchanged (gray).

### Codon usage defines genome-wide clusters and predicts translational linkage to MnmA

To determine whether the translational changes observed in Δ*mnmA* cells are linked to codon usage, we asked whether our TE-defined gene groups shared codon-composition features. Given that codon usage signatures among gene groups are diverse, we used computational approaches to assign codon-defined genes to specific bins. We calculated total codon frequencies (TCF) for each annotated protein-coding gene, using the relative abundance of all 61 sense codons to capture both synonymous codon choice and amino acid demand. Principal component analysis of Z-score–normalized TCF, coupled with Gaussian mixture modeling, resolved the *E. coli* genome into five discrete gene clusters with distinct gene ontology profiles **(Figure 6A, Supplementary Figure S23)**. These clusters showed structured codon signatures rather than random variation of codon assignment **(Figure 6B)**. When codons were grouped by the third-position nucleotide, the clusters segregated clearly into A/U- and G/C-ending codon biases with differences in the use of lysine, glutamine, and glutamic acid codons **(Figures 6C-D)**. MnmA-dependent s²U modifications aid in the decoding of synonymous codons for lysine, glutamine, and glutamate, and we next focused on these codon families. The six codons (AAA/AAG, CAA/CAG, and GAA/GAG) exhibited distinct, cluster-specific enrichment patterns **(Figure 6D)**. Mapping TE-defined genes onto this codon framework demonstrated that all clusters are dysregulated to varying degrees, highlighting the global effect that removal of MnmA has on translation. The most significantly MnmA-sensitive clusters, where all quadrants of translation were affected, were cluster 2 (GAA, GAG, CAA, and CAG leaning) and cluster 1 (AAA, AAG, GAA, GAG, CAA, and CAG leaning) **(Figure 6E)**. Notably, the regulators *rpoS, oxyR,* and *fliA* are all found in Cluster 2. Additional analysis of the TE-up and TE-down genes across genotype and treatment comparisons revealed condition-specific codon-linked patterns **(Supplementary Figures S24–S27)**.

**Figure 6.**
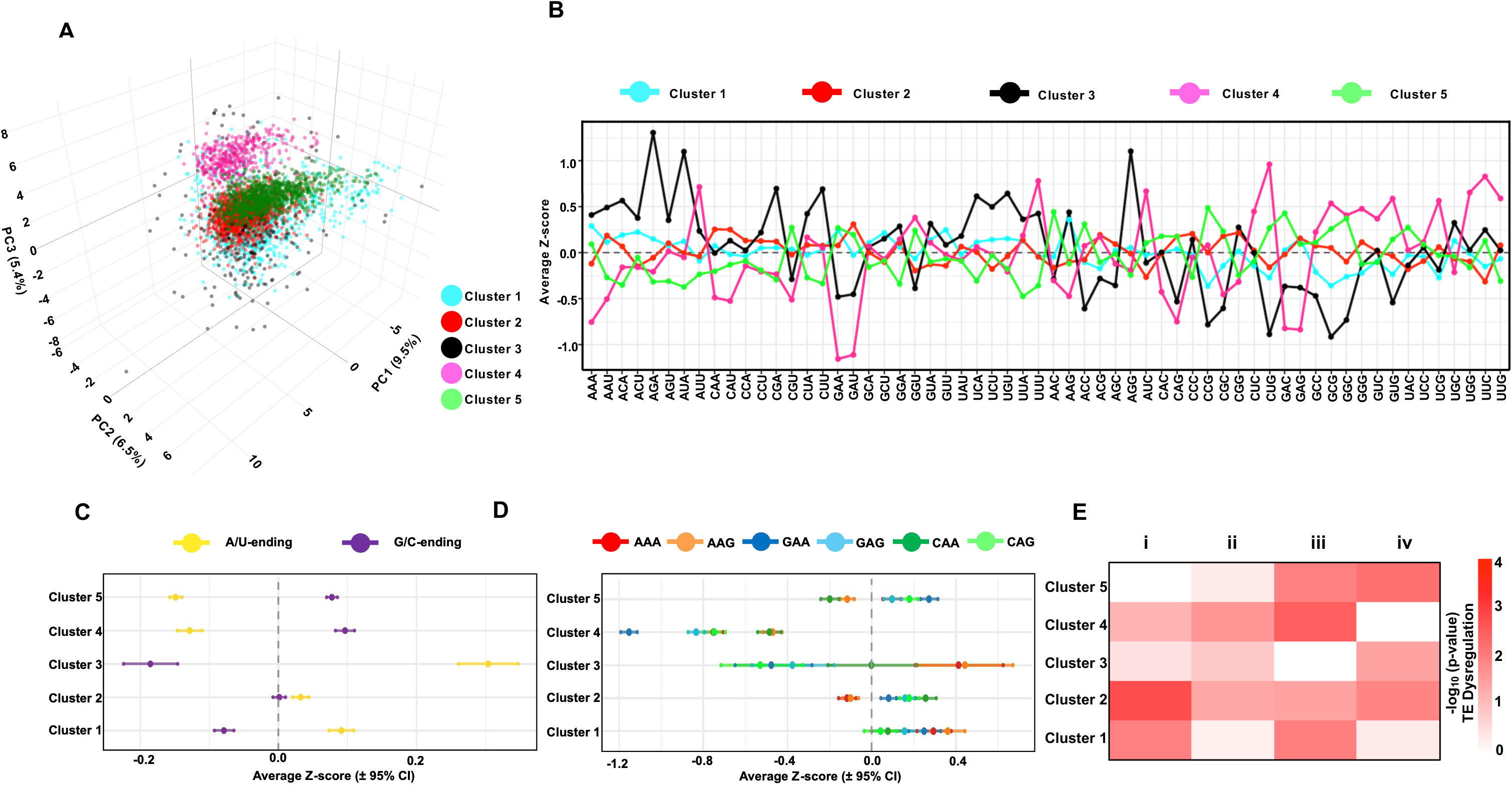
Genome-wide codon usage clustering links patterned groups to MnmA. **(A)** PCA of Z-score normalized TCF across all protein-coding genes in *E. coli*. Genes were clustered using a Gaussian mixture model (GMM), and model selection was based on the Bayesian Information Criterion (BIC). Each point represents a gene colored by its assigned codon-usage cluster. **(B)** Cluster-specific codon signatures showing mean Z-scores for each of the 61 sense codons across the five TCF-derived clusters. **(C)** Mean codon Z-scores ±95% confidence intervals for A/U-ending (yellow) and G/C-ending (purple) codons across the 5 clusters. Positive values indicate enrichment relative to genome averages, whereas negative values indicate depletion. **(D)** Cluster-specific signatures for Lys, Glu, and Gln codon pairs affected by MnmA-dependent wobble U34 chemistry: AAA/AAG, GAA/GAG, and CAA/CAG. Values are shown as mean Z-scores ± 95% CI. **(E)** Heatmap showing the significance of overrepresentation for each cluster within four translational classes for Δ*mnmA*-untreated vs WT-untreated: i = TE_UP_mRNA_DW, ii = TE_DW_mRNA_DW, iii = TE_DW_mRNA_UP, and iv = TE_UP_mRNA_UP. Values represent −log₁₀(p-value) from Fisher’s exact tests, with higher values indicating stronger enrichment.

## Discussion

### Wobble uridine modifications couple codon decoding to bacterial fitness

Wobble uridine modifications provide a conserved layer of translational regulation that helps bacteria match codon-decoding capacity to physiological demand. In Gram-negative cells like *E. coli,* MnmA-dependent s^2^U formation at U34 is important for the decoding of A-ending codons for lysine (AAA), glutamine (CAA), and glutamic acid (GAA) (32,58,61–64). Defects in these modifications have been linked to impaired growth, altered cell physiology, and translational deficiencies (35,38,59). Our findings extend this framework by showing that loss of MnmA depletes s²U-dependent wobble modifications and compromises bacterial growth, oxidative stress defense, and protein output of key stress-response regulators. More broadly, these results support a model in which wobble U34 chemistry not only maintains decoding fidelity but also helps prioritize the translation of codon-defined gene programs required for bacterial fitness.

### Loss of *mnmA* uncouples transcriptional activation from protein output

A central finding of this study is that MnmA-deficient cells mount a broad adaptive transcriptional response but fail to translate it into appropriate protein accumulation. Previous studies have shown that stress can dynamically remodel tRNA modifications to tune translation (11,13,65). However, our data indicate that in *E. coli,* s^2^U abundance is not dynamically reprogrammed. In contrast, under the conditions tested here, *E. coli* s²U abundance did not show evidence of dynamic stress-induced reprogramming. Instead, MnmA-dependent thiolation appears to define a constitutive decoding capacity that varies with growth state and is required when translational demand increases. Loss of MnmA eliminates s²U-dependent wobble modifications, including downstream ges²U and se²U derivatives, thereby imposing a persistent translational limitation that becomes more severe during oxidative stress. It also compromises the ability of the ges²U and se²U derivatives to support the translation of G-ending codons. Within this framework, Δ*mnmA* cells appear to compensate transcriptionally for the underlying translational limitation. The Δ*mnmA* cells show broad mRNA reprogramming, including basal activation of stress-linked regulons, consistent with an attempt to adapt to impaired proteome production, but fail to accumulate key regulatory and detoxification proteins. Polysome profiling further supports this model **(Figure 7)** and shows that oxidative stress exacerbates an existing translational defect in Δ*mnmA* cells. Thus, MnmA loss results in a dramatically altered state, leading to sensitivity to H_2_O_2_-induced oxidative stress.

**Figure 7.**
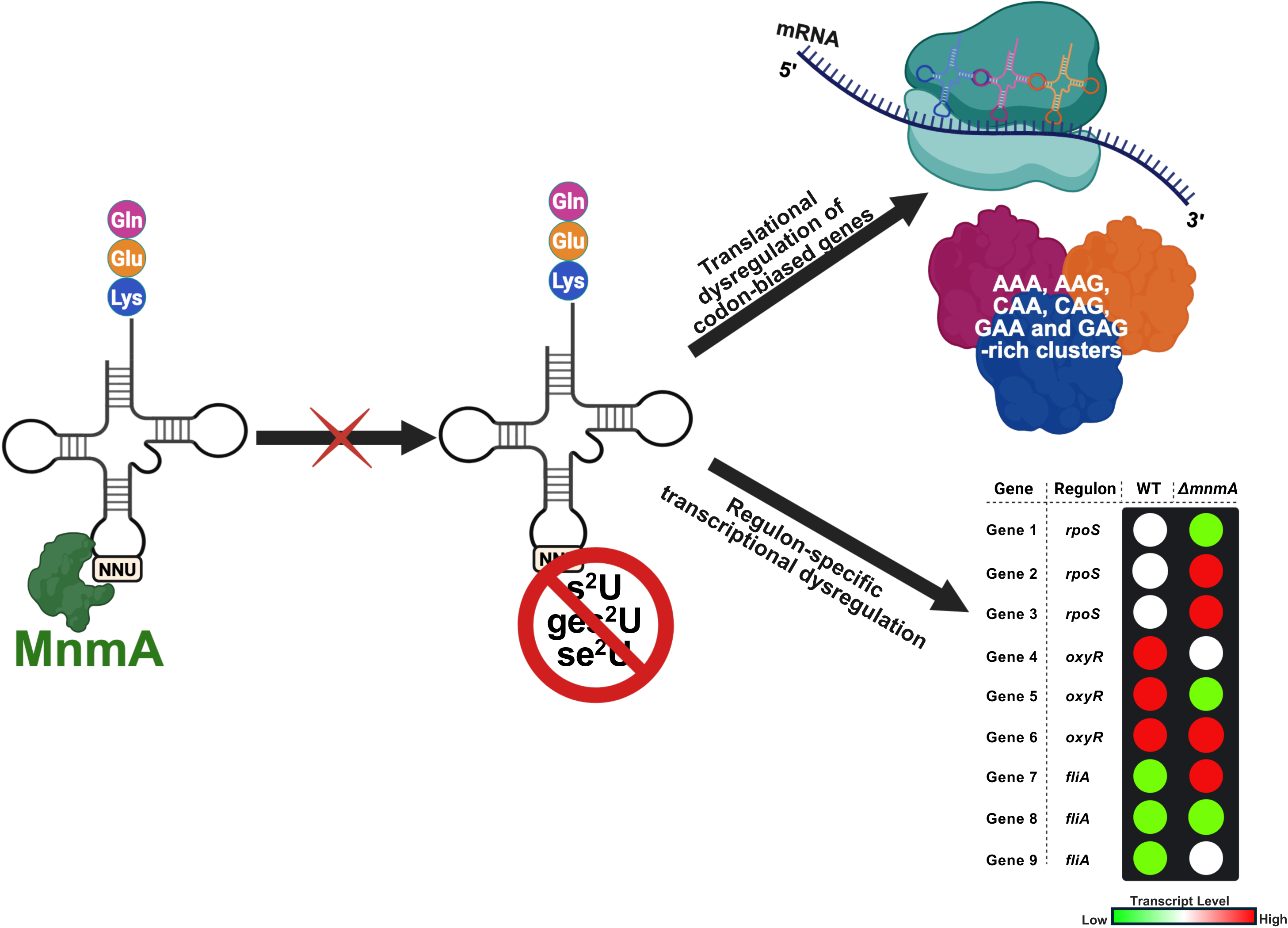
Working model linking MnmA-dependent wobble-U chemistry to codon-biased translation and regulon-level transcriptional remodeling. MnmA catalyzes s²U formation at U34 on tRNAs that decode Lys, Glu, and Gln codons. Loss of MnmA prevents s²U formation and blocks downstream wobble-U remodeling to ges²U and se²U, disrupting the modification pathway on these tRNAs. This defect in MnmA-linked tRNA modification is proposed to affect gene expression through two connected outputs: selective translational dysregulation of codon-biased gene clusters enriched for AAA/AAG, CAA/CAG, and GAA/GAG codons, and regulon-specific transcriptional dysregulation involving *rpoS*-, *oxyR*-, and *fliA*-associated gene programs. Together, this model links MnmA-dependent wobble-U chemistry to selective protein output and genome-wide regulatory imbalance across stress- and other regulatory transcriptional programs.

### MnmA supports cellular adaptation by maintaining protein output from key regulatory factors

MnmA-catalyzed s^2^U modification has been linked to cellular adaptation (58), protein synthesis, and increased cellular growth (66). Our results extend this role by showing that MnmA deficiency reduces protein accumulation of multiple regulatory factors, including RpoS, OxyR, and FliA. RpoS is a master regulator of stationary-phase adaptation (67) and oxidative stress resistance (68,69), and its loss in *ΔmnmA* cells under both basal and H₂O₂-treated conditions suggests that MnmA-dependent decoding capacity is required for RpoS protein accumulation across growth and stress states. Similarly, reduced OxyR and FliA protein levels indicate that MnmA loss affects additional regulatory circuits that control oxidative stress defense and flagellar/motility programs. Studies have shown that OxyR is induced under oxidative stress (70–72), but our work supports the view that OxyR is also basally expressed. Although prior work has emphasized stress-induced remodeling of tRNA modifications, our findings suggest that MnmA-dependent s²U provides a constitutive translational capacity that becomes especially important when cells must rapidly adjust regulatory protein output. Thus, MnmA loss compromises adaptation not by eliminating transcriptional responses, but by limiting production of the regulatory proteins needed to coordinate stress defense, growth-state transitions, and motility-associated programs.

### TCF-based codon clustering reveals MnmA-linked gene programs in *E. coli*

A key conceptual advance of this work is linking MnmA-dependent wobble-U chemistry to genome-wide codon organization. Codon adaptation influences both mRNA stability and translation (73), and systems-level studies in eukaryotes have shown that distinct gene classes can differ in their dependence on tRNA pools and tRNA modifications (60,62). Here, we extended this framework to *E. coli* by showing that total codon frequency analysis stratifies the genome into five codon-defined gene groups with distinct functional enrichments. While all clusters are affected, the most severe dysregulation in the MnmA-deficient background was observed in clusters with specific profiles of Gln (CAA, CAG) and Glu (GAA, GAG) codons, with a lesser contribution for Lys (AAA, AAG. These findings suggest that codon composition helps define which gene programs are most vulnerable to loss of MnmA-dependent modifications, with the limitations discussed below being relevant. In particular, the regulatory nodes in oxidative and general stress pathways, including *rpoS, oxyR,* and *fliA*, occupy distinct codon-usage space (Cluster 2) associated with MnmA. Thus, rather than functioning solely as a facilitator of individual tRNA–codon interactions, MnmA operates within a broader regulatory architecture that promotes the translation of many Gln, Glu and Lys codon-defined gene programs central to regulation and stress adaptation.

### Study limitations

While our data provide important details on gene expression, there are limitations on the analysis and interpretation of MnmA effects. One technical limitation is that we used tagged proteins to study several stress-response genes because suitable native antibodies were not available. Although these tagged constructs allowed us to compare protein levels, plasmid-based expression does not fully reflect natural gene expression. Tag placement and differences in the 5′ and 3′ gene context may also affect regulation. The translation data should also be interpreted with caution. Adaptive transcriptional changes in Δ*mnmA* cells complicate codon-specific analyses. In bacteria, transcription and translation are coupled, so changes in mRNA abundance at the gene and operon levels can alter which transcripts are translated. This makes codon-based analysis complicated, with codon usage tied to the changing gene-expression needs of Δ*mnmA* cells. TE data also have limitations because they measure polysome occupancy, which reflects mRNA loading onto ribosomes as well as pausing and elongation. Polysome-bound mRNAs can include single genes or operons, and genes near the 5′ end of an operon can influence the polysome signals of downstream genes. Thus, codon-enrichment patterns in Δ*mnmA* cells likely result from several combined effects, including altered decoding capacity, compensatory transcriptional adaptation, and operon structure. Ribosome profiling would help distinguish impaired initiation, altered elongation, and changes in mRNA loading for specific genes and codons. Overall, our codon-clustering analysis shows links between codon use, translation patterns, and MnmA sensitivity, but it does not prove direct cause-and-effect relationships for individual genes. Future studies using synonymous recoding, native-locus reporter constructs, and targeted rescue of tRNA modification states will be needed to test whether specific codon patterns are enough to cause MnmA-dependent translational control.

## Data availability

The data underlying this article are available in the article and in its online supplementary material.

For high-throughput data:

RNA-seq raw data (FASTQ files) and processed data have been deposited in NCBI’s Gene Expression Omnibus (GEO) and are accessible through the GEO accession numbers GSE330173 and GSE330174 (currently under embargo).

RNA-seq and codon clustering scripts are available at https://github.com/HO24-R/MnmA_manuscript

Custom plasmids/strains are available upon request.

## Acknowledgments

We are grateful to members of the Begley, Sheng, and Dedon labs for providing input and helpful advice. The authors are grateful for support from the National Institutes of Health (ES026856, ES031529, GM070641, CA274603) and the National Research Foundation of Singapore through the Singapore-MIT Alliance for Research and Technology Antimicrobial Resistance Interdisciplinary Research Group. Pictorial graphics were generated using Inkscape and BioRender.

## AI use statement

After compiling the figures and drafting the manuscript, the authors used OpenAI’s ChatGPT 5.5 (CHO) and Microsoft Copilot (TJB) under a paid premium license to rephrase parts of the manuscript for clarity, conciseness, and grammatical correctness. In addition, OpenAI’s ChatGPT 5.5 was used to help facilitate and refine parts of the R scripts. After using this tool/service, the author(s) reviewed and edited the contents and results as needed and take(s) full responsibility for the content of the publication.

## Author contributions and ORCID details

**HCO:** Conceptualization, investigation, graphical abstract, methodology, validation, data curation, formal analysis, visualization, writing—original draft, and writing—review and editing. **DE:** Data analysis, visualization, and writing—review and editing. **CM:** Graphical abstract, methodology, and writing—review and editing. **CR:** Methodology and writing—review and editing. **KU:** Methodology. **ETD:** Methodology and writing—review and editing. **RA:** Data analysis. **AD:** Methodology **QL:** Methodology and writing—review and editing. **PCD:** Conceptualization, supervision, validation, and writing—review, and editing. **JS:** Conceptualization, supervision, validation, and writing—review and editing. **TJB:** Conceptualization, formal analysis, supervision, validation, visualization, resources, funding acquisition, and writing—review and editing.

