## Supplemental Figures for "The Wobble Uridine tRNA Writer MnmA Shapes Codon-Dependent Stress Response Systems"

### Figure S1

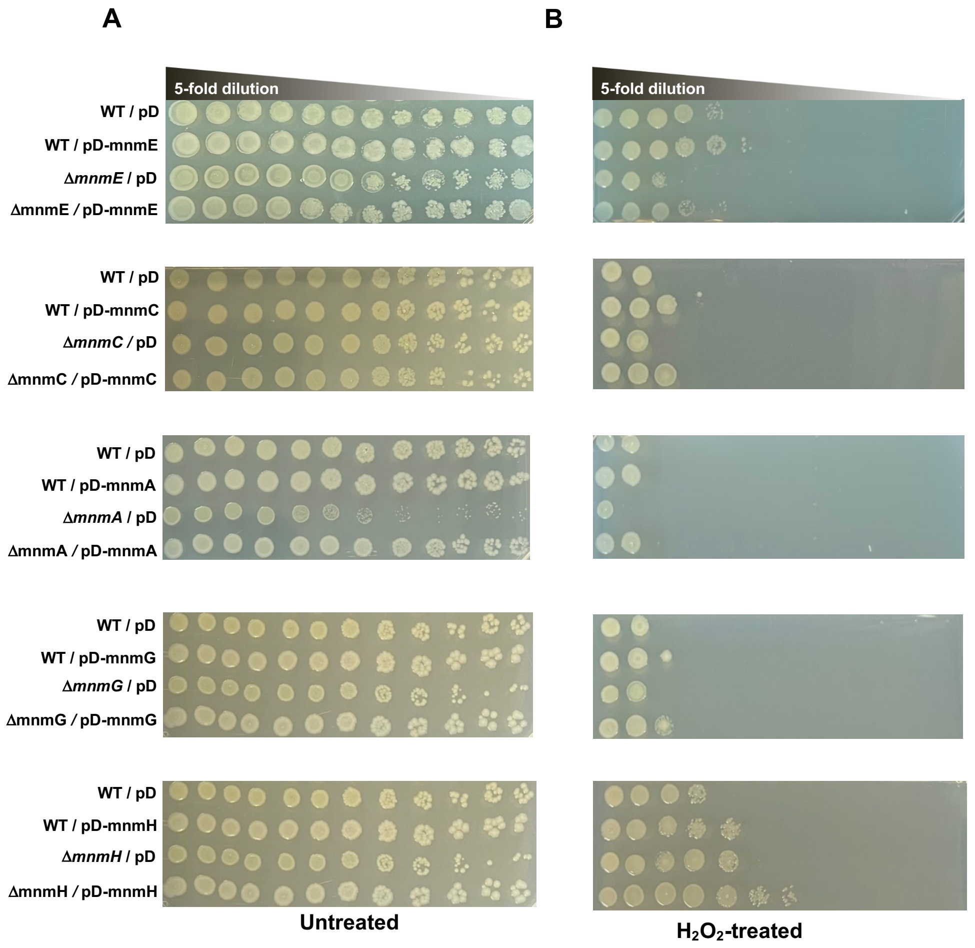


**Supplementary Figure S1: All mutants +/- $H_2O_2$. All mutants +/- $H_2O_2$ and CAM.** The growth sensitivity assay was performed on Δ*mnmA*, Δ*mnmC*, Δ*mnmE*, Δ*mnmG*, and Δ*mnmH*, with the left panel (A) untreated and the right panel (B) treated with 5 mM $H_2O_2$.

Figure S2

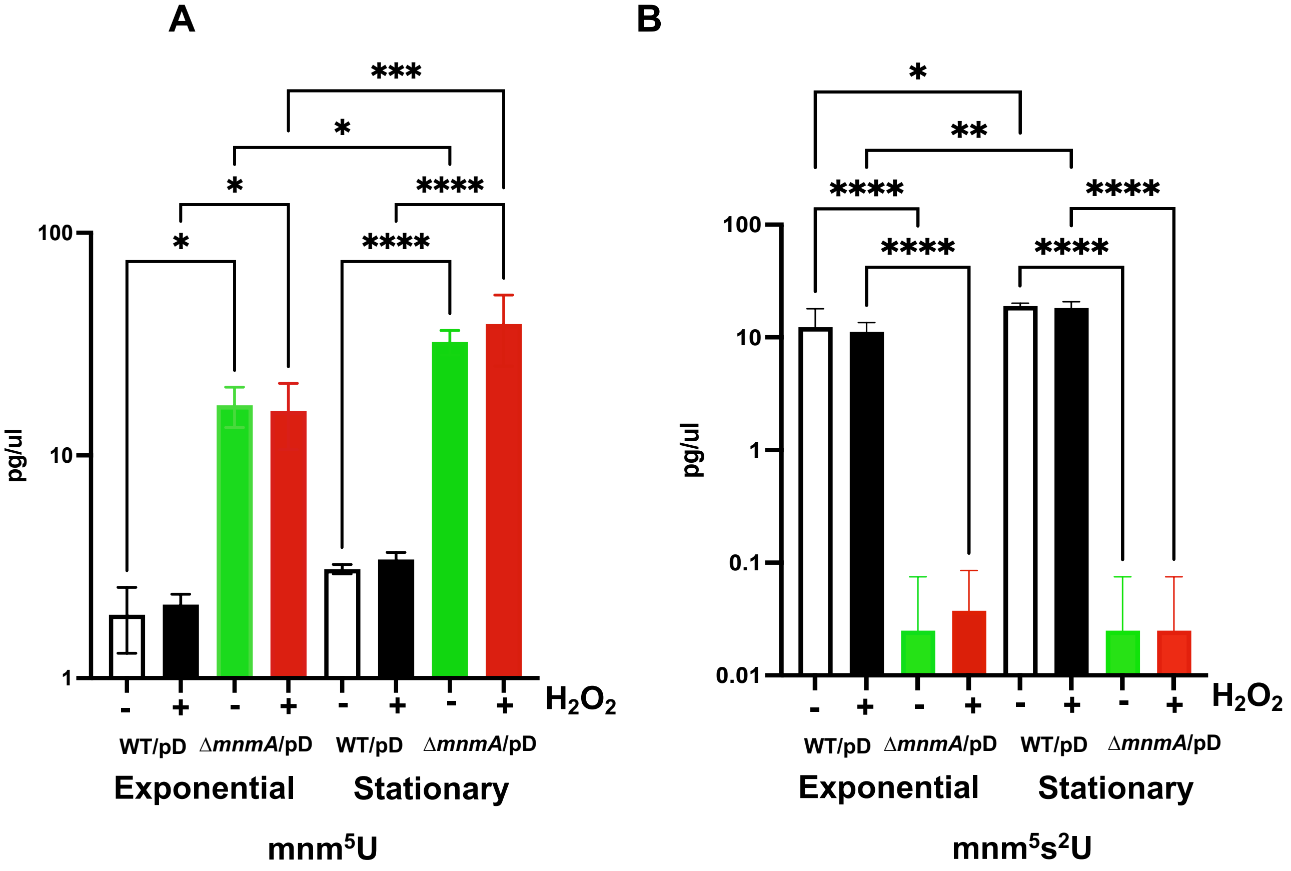


**Supplementary Figure S2: *E. coli* tRNA modifications dynamics in exponential and stationary growth phases.** Growth-state-dependent levels of wobble $U_{34}$ tRNA modifications require MnmA. (A) $mnm^5U$, and (B) $mnm^5s^2U$ levels were detected by LC-MS/MS in total tRNA isolated from *E. coli* WT and Δ*mnmA* cells during exponential (exp) and stationary (sta) phases, with or without $H_2O_2$ treatment (tr). Data was analyzed using ANOVA and represents mean ± SD; n=3. **p<0.05, **p<0.01, ***p<0.001, ****p<0.0001*

**Figure S3**

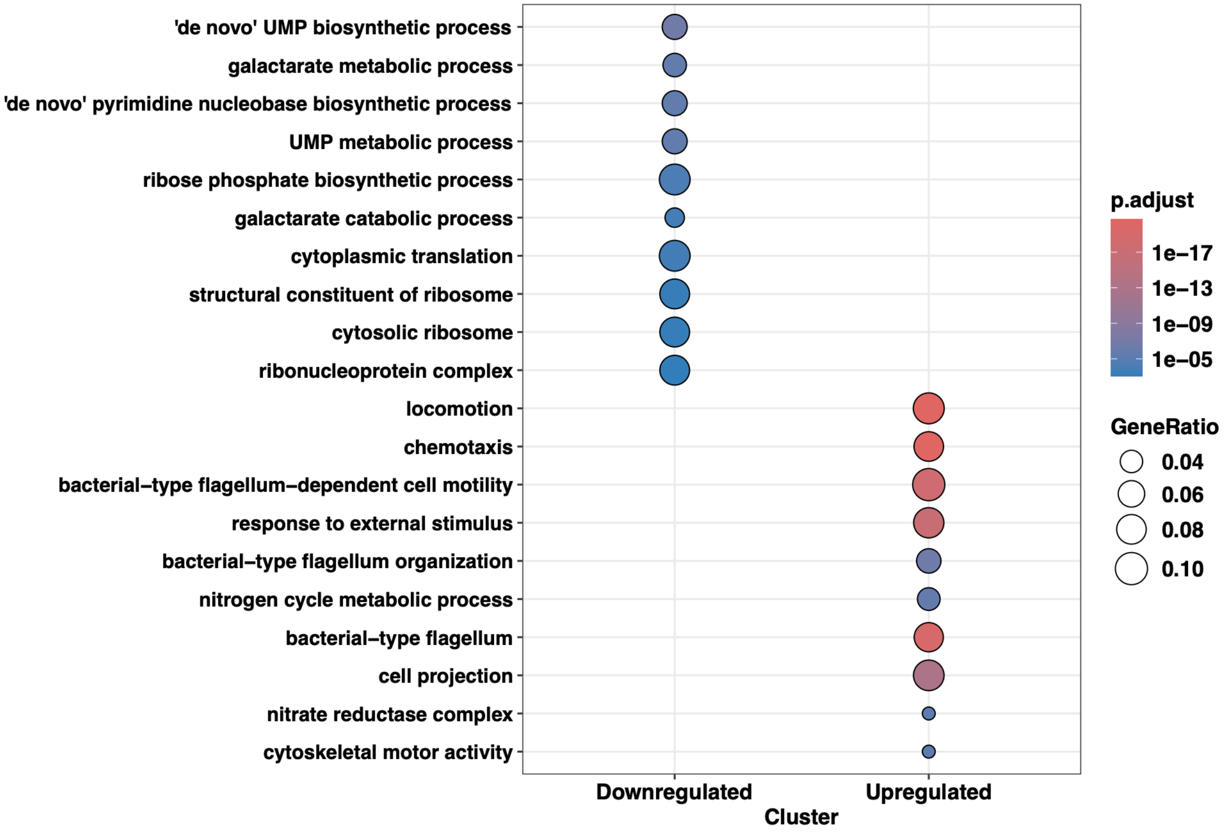


**Supplementary Figure S3: Gene ontology analysis for MnmA-deficient and WT cells under basal conditions.** Total RNA from Δ*mnmA* untreated and WT untreated cells at the exponential phase was purified and analyzed by NGS mRNA-seq (n = 4). Differentially expressed genes were defined as |log2FC| ≥ 1.50 and padj ≤ 0.05, and Gene Ontology enrichment was performed with ClusterProfiler for the downregulated and upregulated gene sets separately. Dot size indicates GeneRatio, and dot color indicates adjusted P value (p.adjust).

**Figure S4**

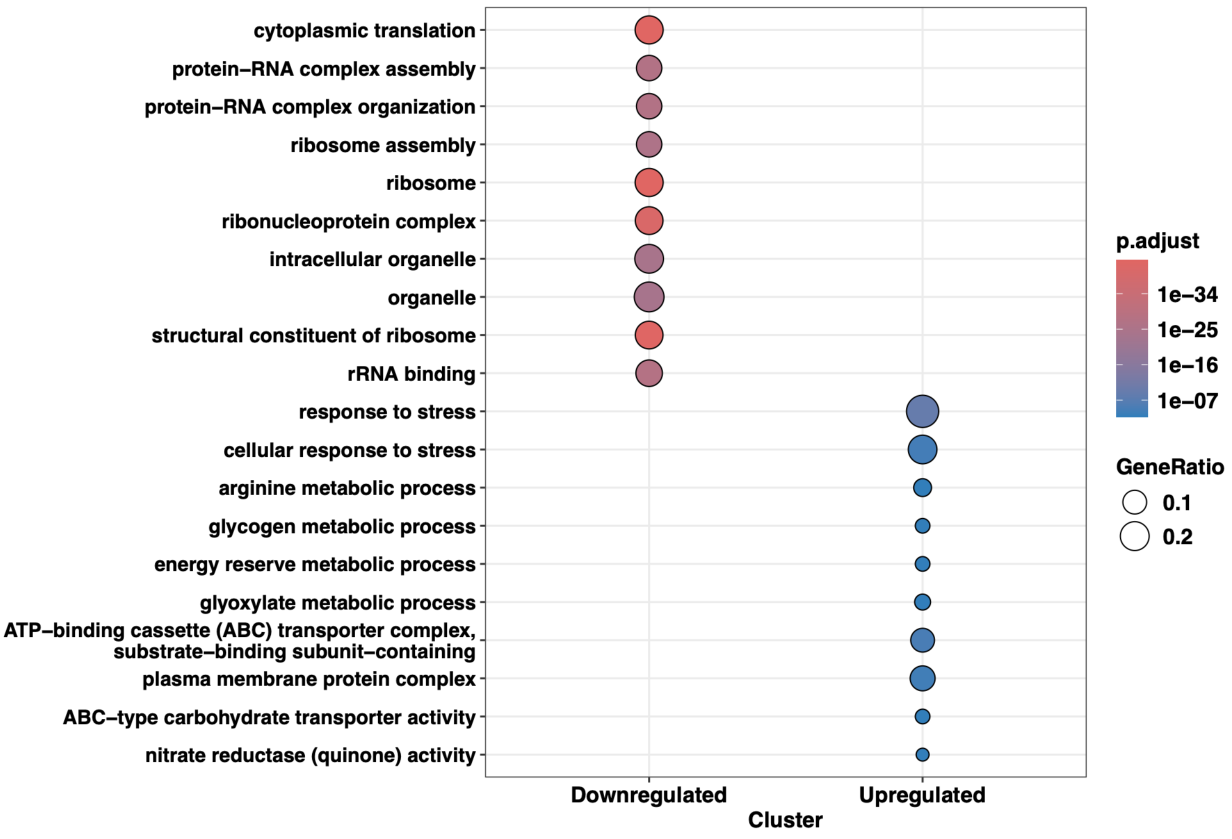


**Supplementary Figure S4: Gene ontology analysis for WT cells under different conditions.** Total RNA from WT-treated and WT-untreated cells at the exponential phase was purified and analyzed by NGS mRNA-seq (n = 4). Genes meeting |log2FC| ≥ 1.50 and padj ≤ 0.05 were split into downregulated and upregulated sets. Gene ontology enrichment was computed using the ClusterProfiler package. Enriched terms are shown as dot plots, with dot size representing GeneRatio and dot color representing p-adjusted values

**Figure S5**

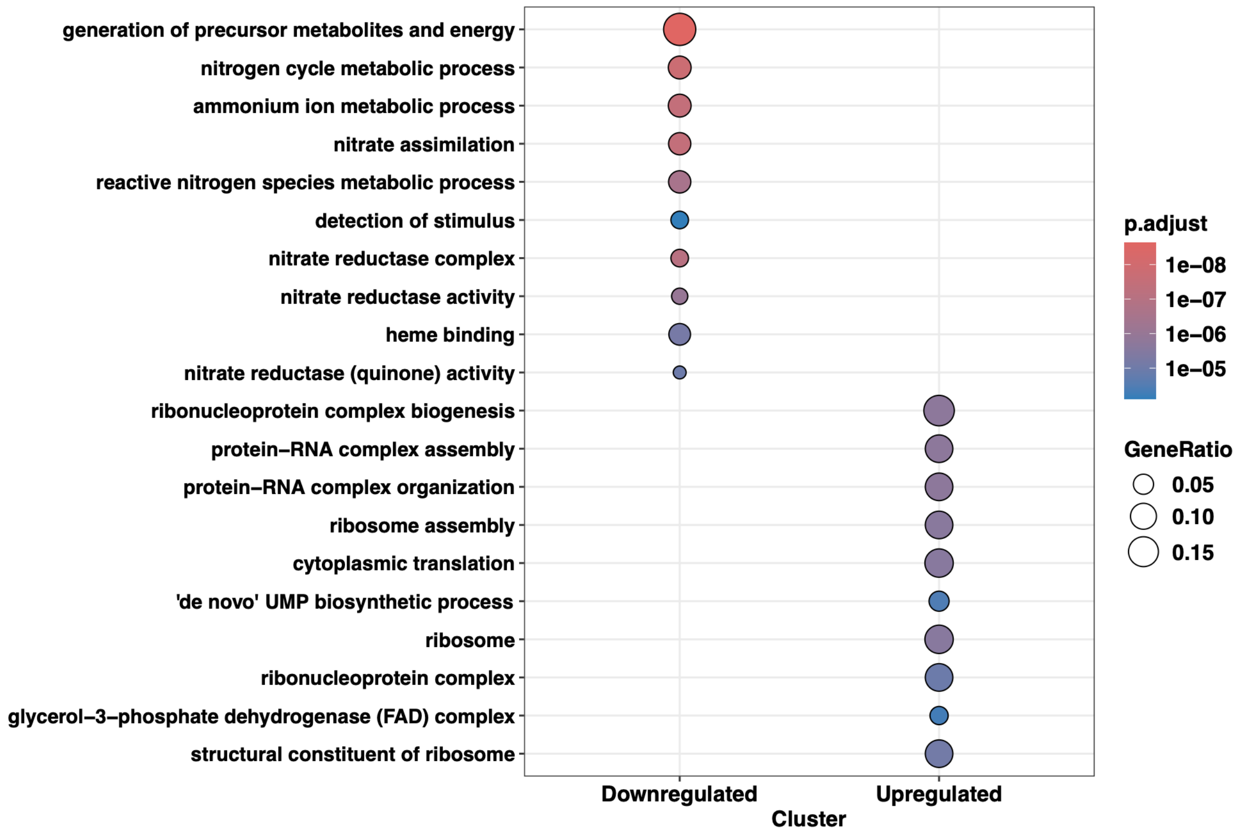


**Supplementary Figure S5: Gene ontology analysis for MnmA-deficient cells under different conditions.** Total RNA from Δ*mnmA*-treated and Δ*mnmA*-untreated cells at the exponential phase was purified and analyzed by NGS mRNA-seq (n = 4). Differential expression cutoffs were |log2FC| ≥ 1.50 and padj ≤ 0.05. clusterProfiler was used to perform gene ontology analysis separately for downregulated and upregulated genes, and the results are displayed using GeneRatio (dot size) and p.adjust (dot color).

**Figure S6**

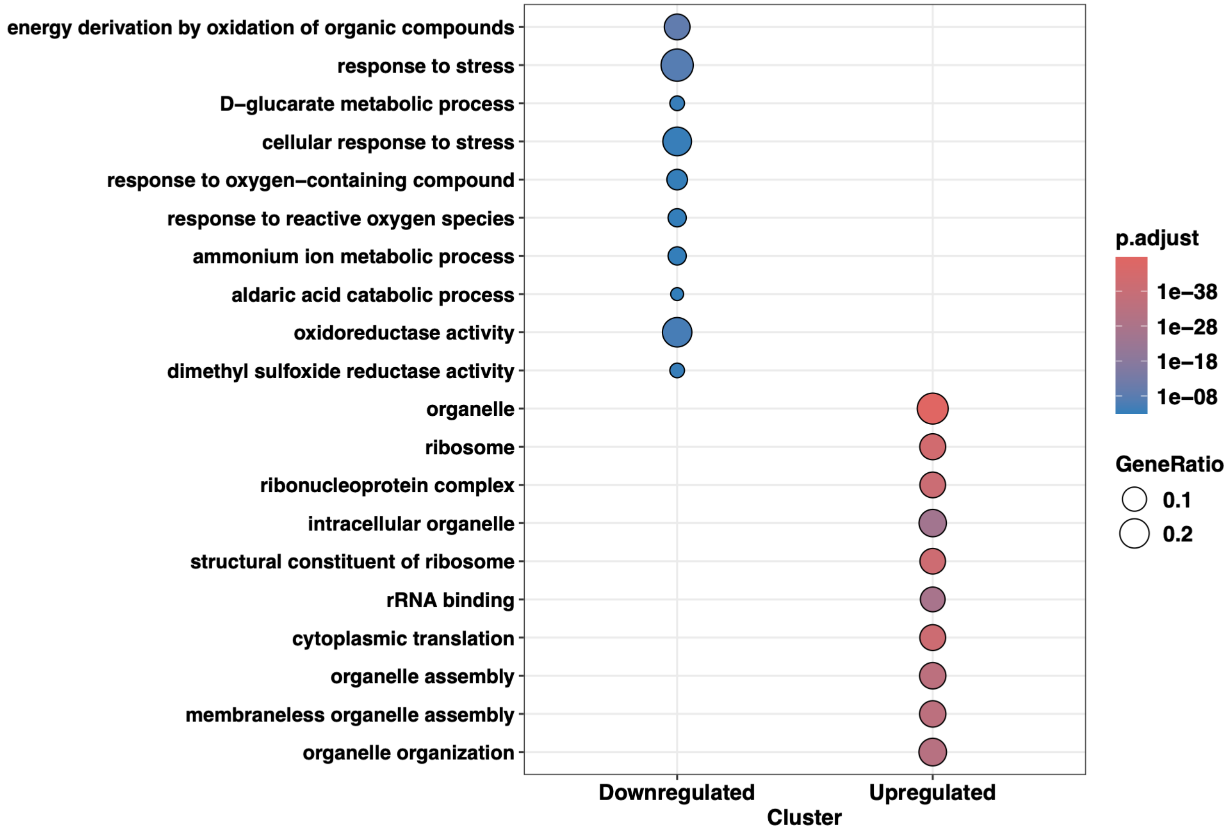


**Supplementary Figure S6: Gene ontology analysis for MnmA-deficient and WT cells under $H_2O_2$-treated conditions.** Total RNA from Δ*mnmA*-treated and WT-treated cells at the exponential phase was purified and analyzed by NGS mRNA-seq (n = 4). Genes were classified as differentially expressed with |log2FC| ≥ 1.50 and padj ≤ 0.05. Gene ontology analysis was performed independently for downregulated and upregulated gene sets using ClusterProfiler. Dot plots display GeneRatio (dot size) and adjusted P value (p.adjust; dot color).

**Figure S7**

**A**

**WT treated vs WT untreated**

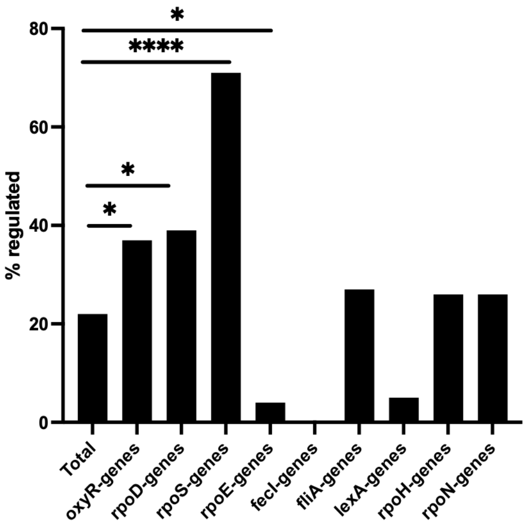


**B**

**Δ*mnmA* treated vs Δ*mnmA* untreated**

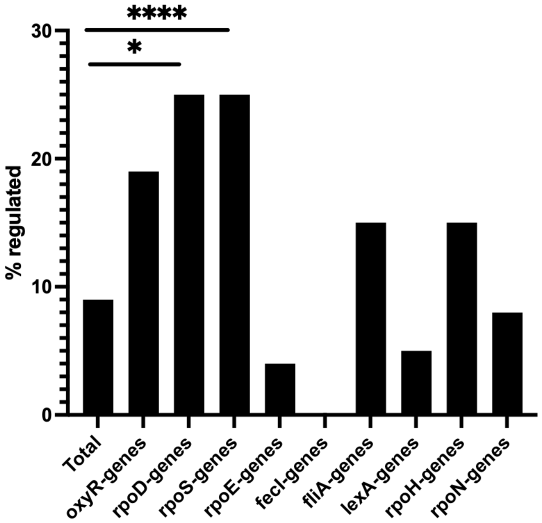


**Supplementary Figure S7: Graph of dysregulated regulons in different group comparisons.** Fischer's test of significance for dysregulated regulons was analyzed at the exponential phase for the *E. coli* regulons and presented as bar graphs for (A) WT treated vs. WT untreated and (B) Δ*mnmA* treated vs. Δ*mnmA* untreated. (*p < 0.05, ****p < 0.0001).

**Figure S8**

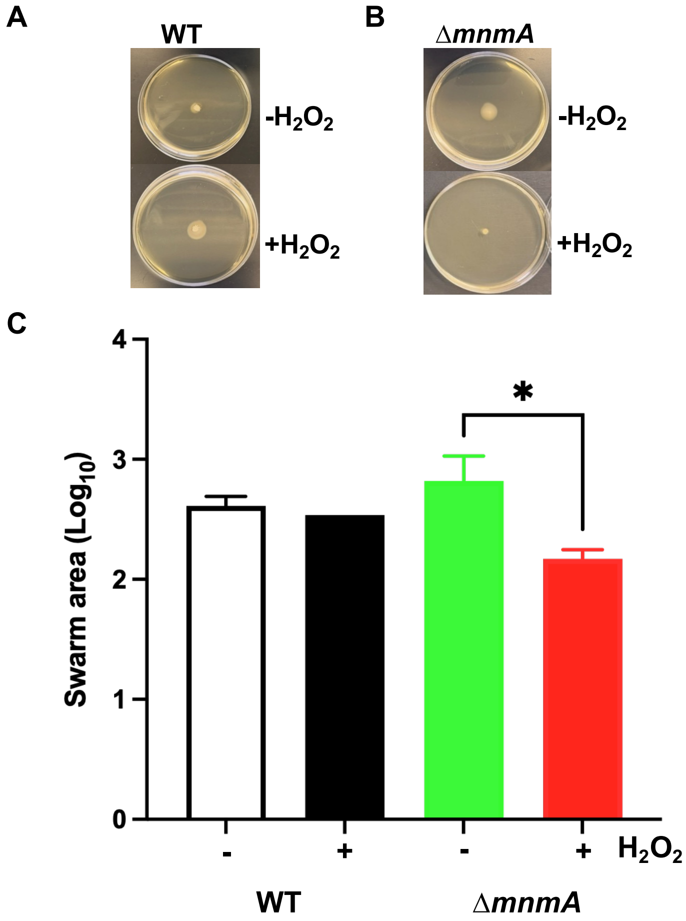


**Supplementary Figure S8: *E. coli* MnmA is necessary for swarming motility.** Swarming motility of *E. coli* WT and Δ*mnmA* on $H_2O_2$-treated and untreated motility agar plates. 3 ul of cells in the exponential growth phase were inoculated on 0.35% (wt./vol) agar and incubated overnight at 37 $^{o}$C, n = 3. Photographs show representative swarm activity of (A) WT and (B) Δ*mnmA* cells under different treatment conditions. (C) Swarm diameter was measured and analyzed using ANOVA. *$p < 0.05$

**Figure S9**

A

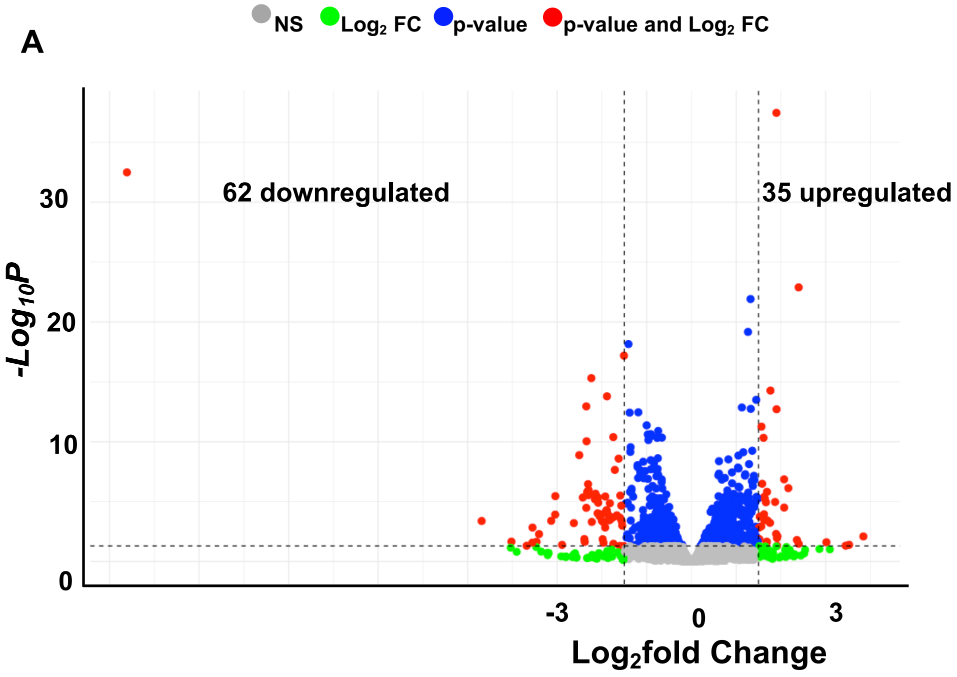


B

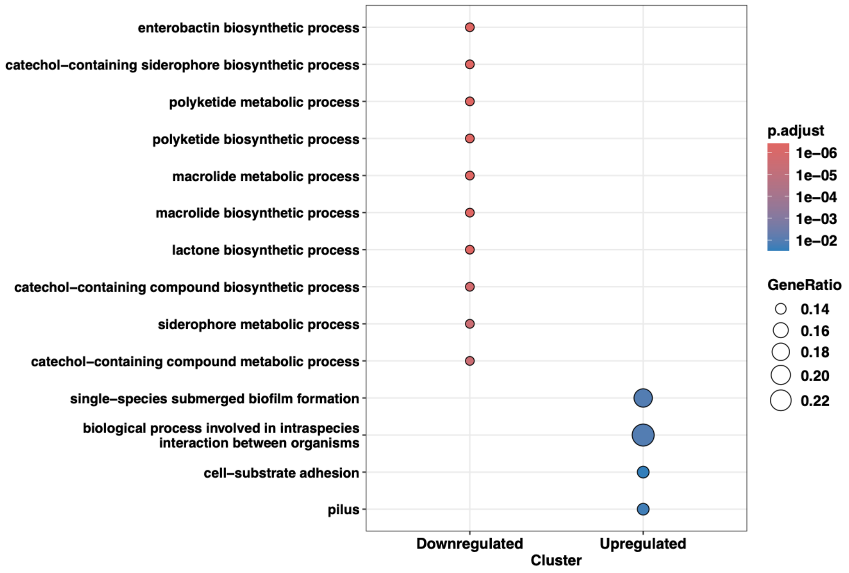


**Supplementary Figure S9: mRNA RNASeq analysis under basal conditions.** Total RNA from Δ*mnmA* untreated and WT untreated cells at the stationary phase was purified and analyzed by NGS mRNA-seq (n = 4). (A) Volcano plot showing differentially upregulated (35) and downregulated (62) genes. Gene ontology analysis was performed using clusterProfiler for (B) downregulated and upregulated genes.

**Figure S10**

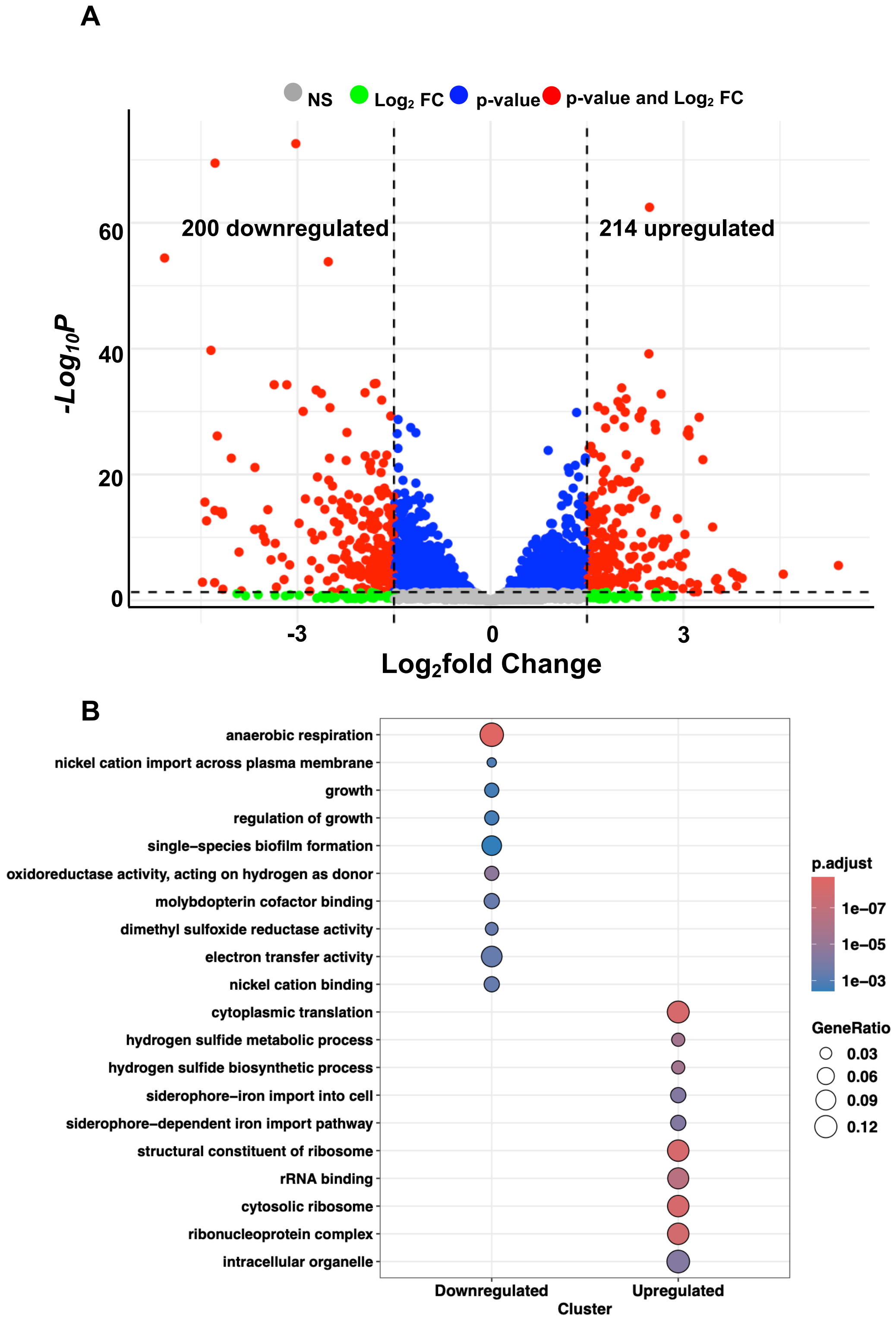


**Supplementary Figure S10: mRNA profile comparison for WT cells.** Total RNA from WT-treated and WT-untreated cells at the stationary phase was purified and analyzed by NGS mRNA-seq (n = 4). (A) Volcano plot showing differentially upregulated (214) and downregulated (200) genes. Gene ontology analysis was performed using clusterProfiler for (B) downregulated genes ($log_2FC \leq 1.50$ and $P_{adj.}$ values $\leq 0.05$), and upregulated genes ($log_2FC \geq 1.50$ and $P_{adj.}$ values $\leq 0.05$).

Figure S11

A

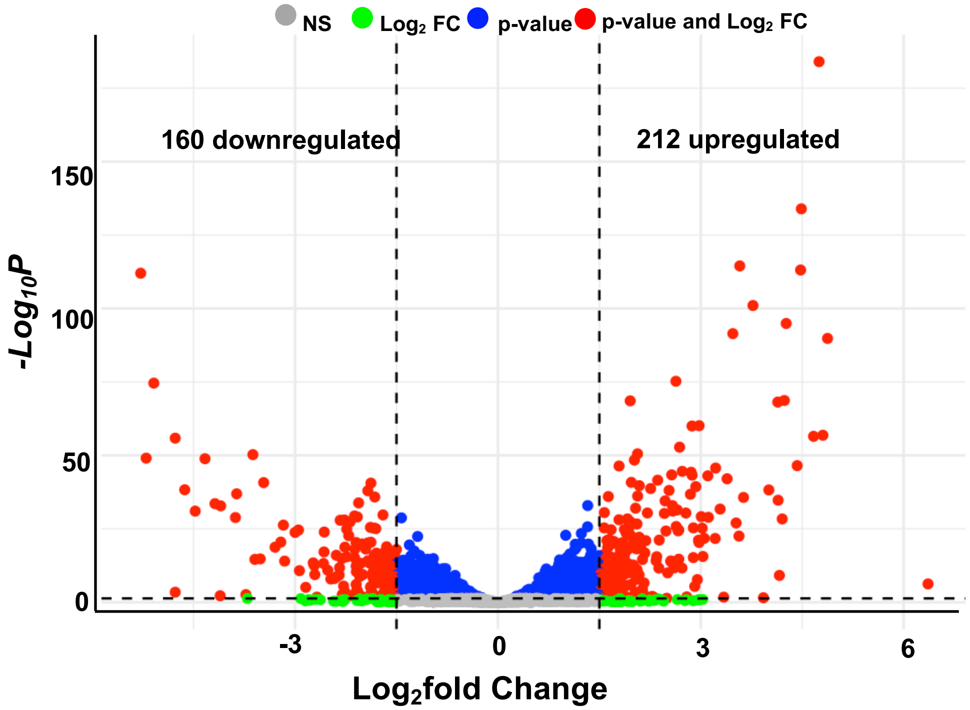


B

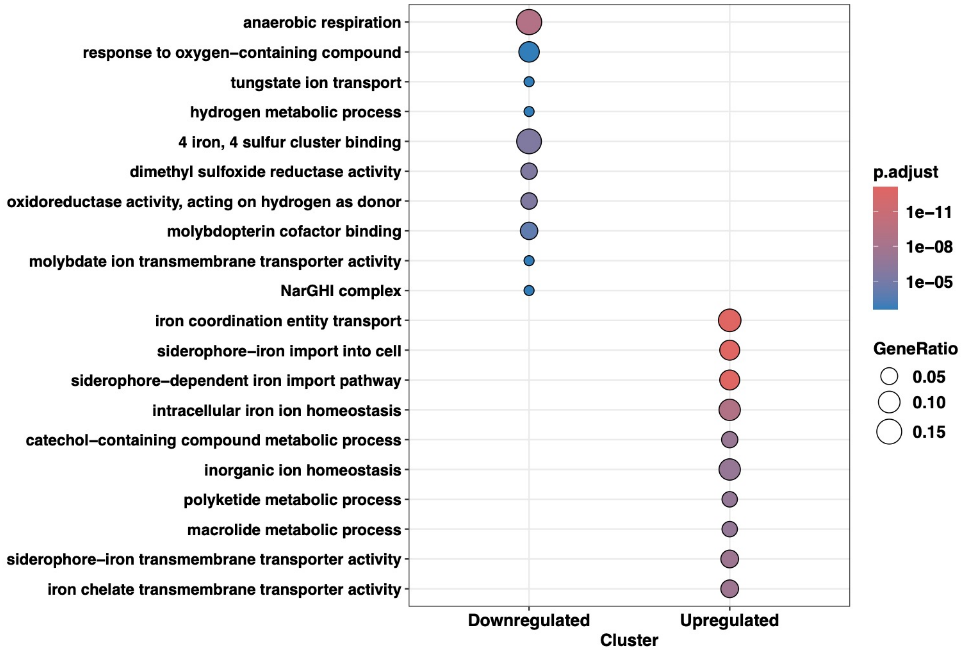


**Supplementary Figure S11: mRNA profile comparison for MnmA-deficient cells.** Total RNA from Δ*mnmA*-treated and Δ*mnmA*-untreated cells at the stationary phase was purified and analyzed by NGS mRNA-seq (n = 4). (A) Volcano plot showing differentially upregulated (212) and downregulated (160) genes. Gene ontology analysis was performed using clusterProfiler for (B) downregulated genes (log2FC ≤ 1.50 and Padj. values ≥ 0.05) and upregulated genes (log2FC ≥ 1.50 and Padj. values ≥ 0.05).

**Figure S12**

**A**

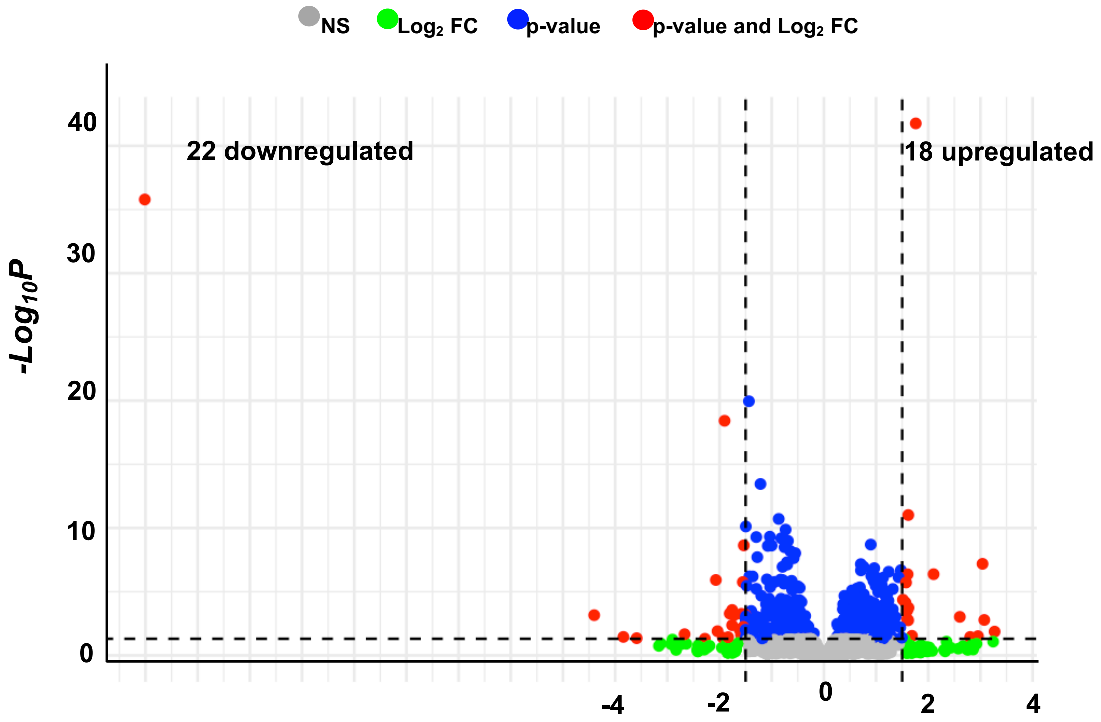


**B**

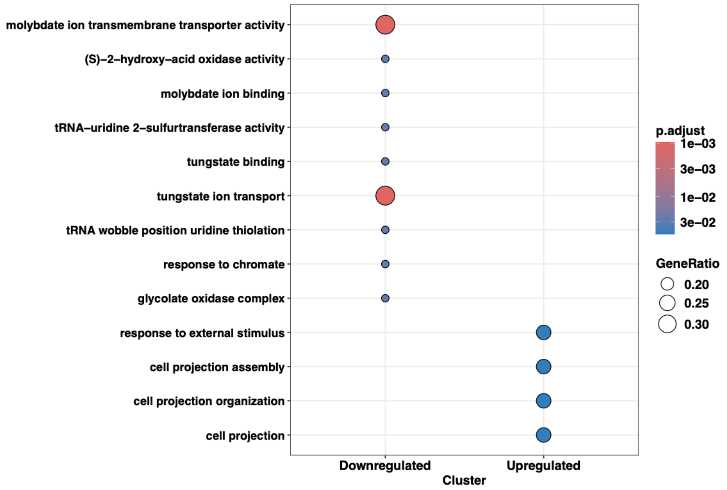


**Supplementary Figure S12: mRNA profile analysis for Δ*mnmA* vs WT cells.** Total RNA from Δ*mnmA*-treated and WT-treated cells at the stationary phase was purified and analyzed by NGS mRNA-seq (n = 4). (A) Volcano plot showing differentially upregulated (18) and downregulated (22) genes. Gene ontology analysis was performed using clusterProfiler for (B) downregulated genes ($log_2FC$ ≤1.50 and $P_{adj.}$ values ≤ 0.05), and upregulated genes ($log_2FC$ ≥ 1.50 and $P_{adj.}$ values ≤ 0.05).

**Figure S13**

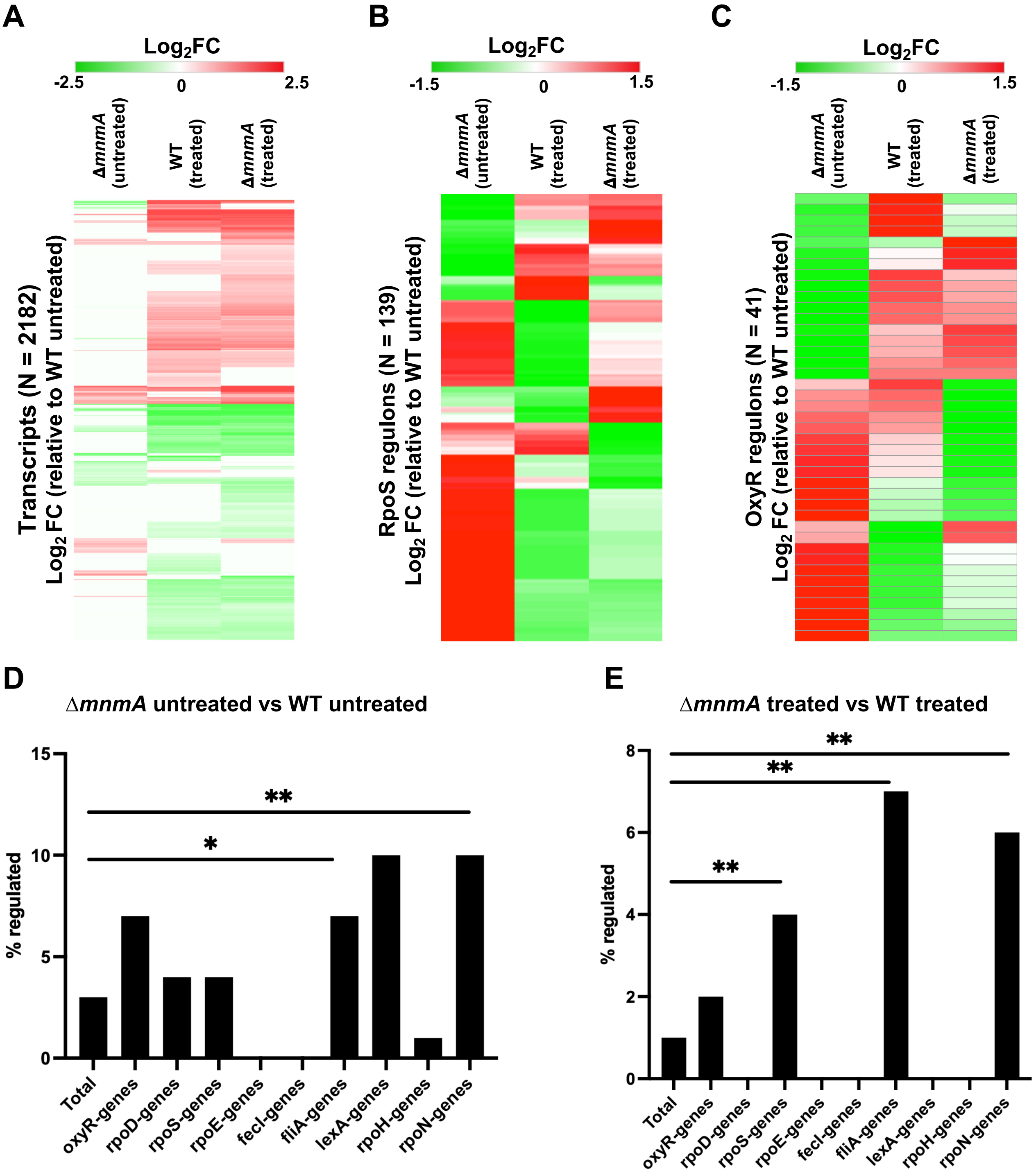


**Supplementary Figure S13: Regulon-specific transcripts are dysregulated in Δ*mnmA* cells during the stationary phase.** Heat map visualizations of clustered genes were analyzed ($log_2FC \geq 1.50$ and $log_2FC \leq 1.50$, $P_{adj.}$ values $\leq 0.05$) for (A) total transcripts regulated (N = 2182), (B) RpoS-linked (N = 139), and (C) OxyR-linked transcripts (N = 41). Fischer's test of significance for dysregulated regulons was analyzed at the exponential phase for the E. coli regulons and presented as bar graphs in (D) Δ*mnmA* untreated vs. WT untreated and (E) Δ*mnmA* treated vs. WT treated. (*p < 0.05, ****p < 0.0001)

### Figure S14

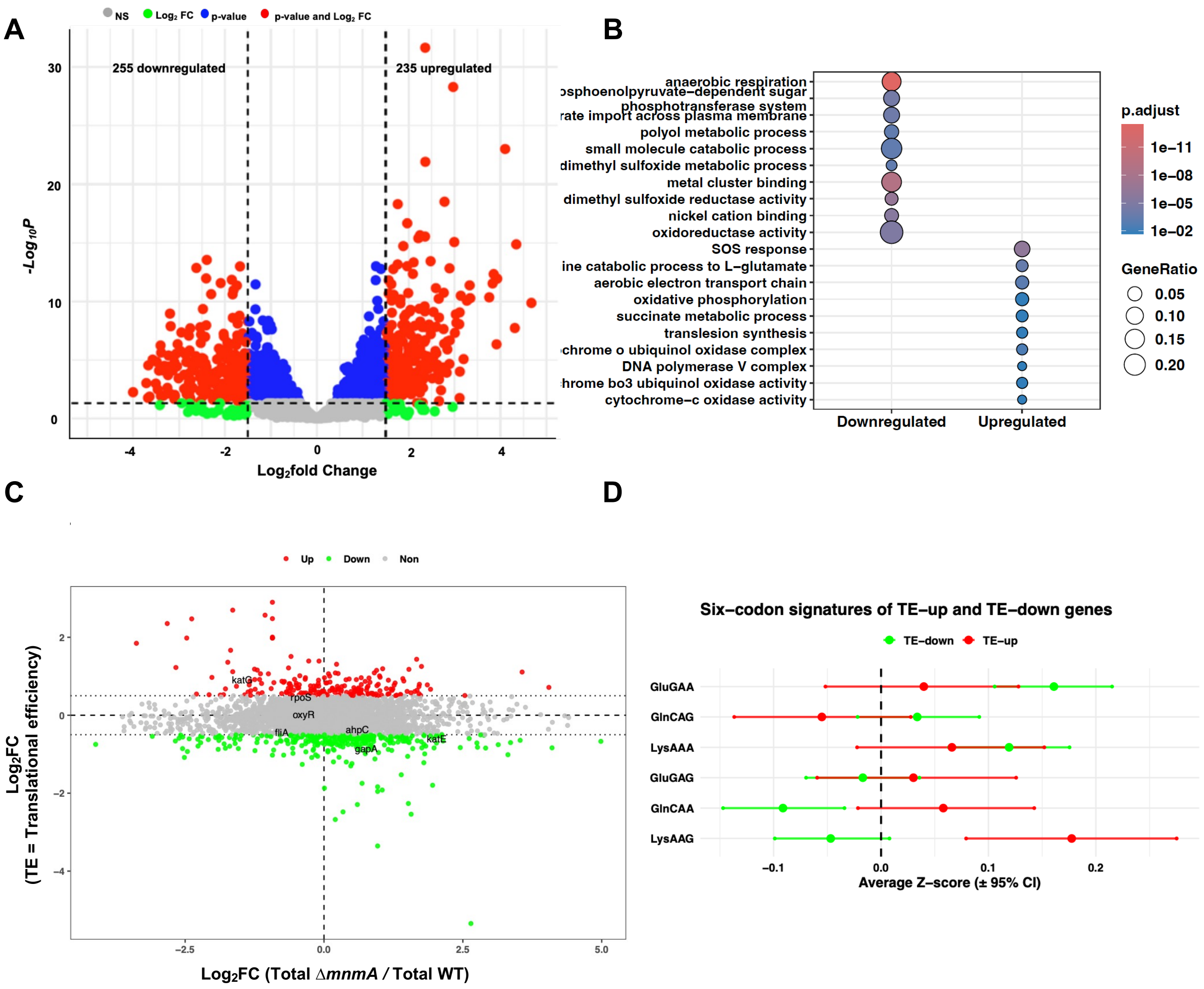


**Supplementary Figure S14. Polysome-associated mRNA profiling of WT cells following oxidative stress exposure.** Total RNA was isolated from polysome fractions of exponentially growing WT cells under untreated and $H_2O_2$-treated conditions, and the samples were analyzed by mRNA-seq ($n$ = 3). (A) Volcano plot showing differentially expressed polysome-associated transcripts in WT-treated cells relative to WT-untreated cells. Significantly upregulated (235) and downregulated (255) genes are indicated. (B) Gene Ontology (GO) enrichment analysis of significantly upregulated and downregulated transcripts using clusterProfiler. Dot size represents the gene ratio, and color indicates the adjusted P-value. (C) Translational efficiency (TE) plotted against total mRNA changes for WT-treated relative to WT-untreated cells. TE was calculated as the ratio of polysome-bound mRNA to total mRNA. Genes are classified as TE-up (red), TE-down (green), or unchanged (gray). Dashed lines indicate the mRNA and TE thresholds used for classification. Selected genes are labeled. (D) Six-codon signatures of TE-up and TE-down genes in WT cells following treatment. Average codon Z-scores (±95% CI) are shown for AAA/AAG, GAA/GAG, and CAA/CAG. TE-up genes are shown in red and TE-down genes in green. The dashed vertical line marks the genome-wide average codon usage after normalization.

### Figure S15

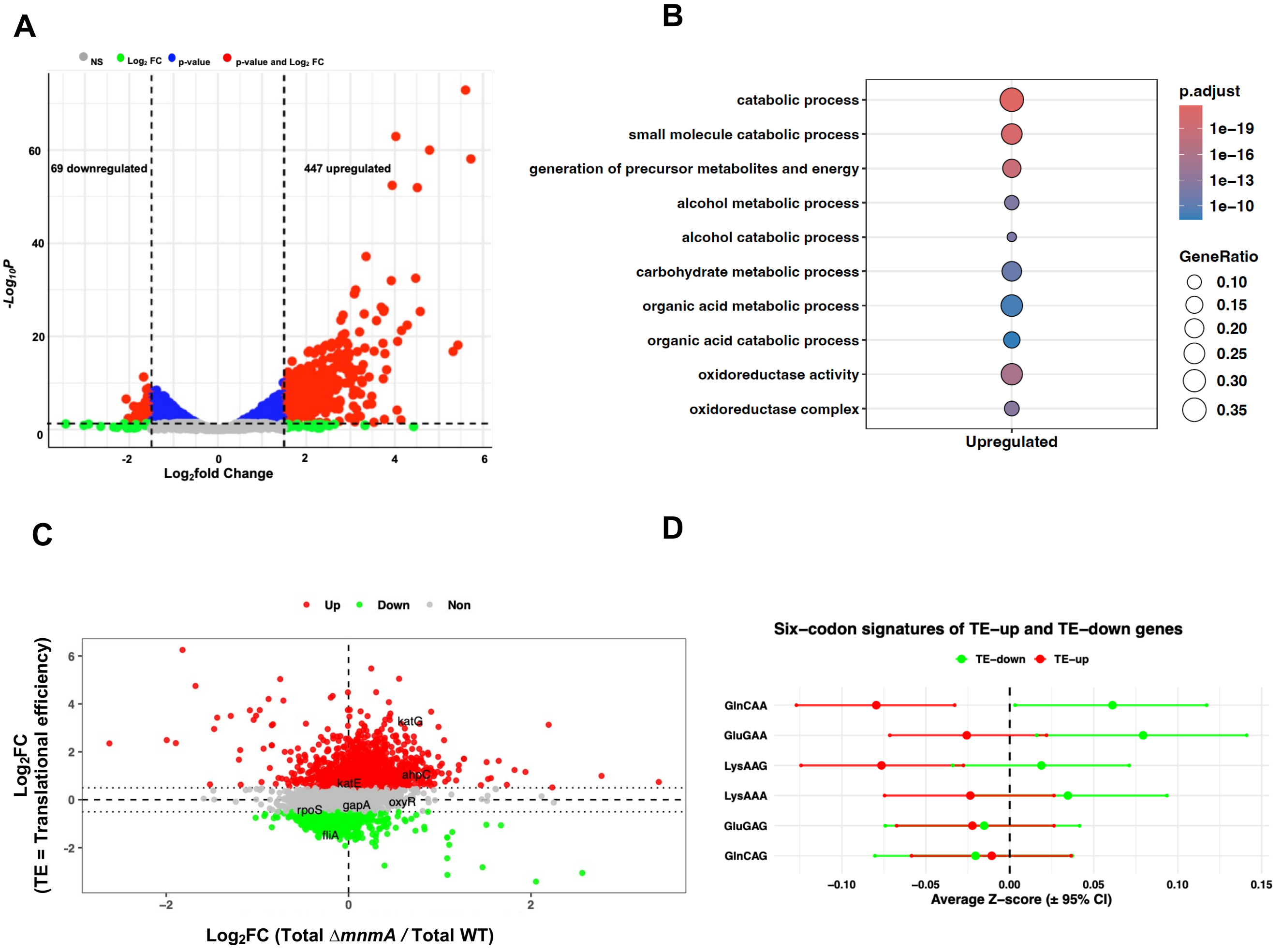


**Supplementary Figure S15. Polysome-associated mRNA profiling of Δ*mnmA* cells following oxidative stress exposure.** Total RNA was isolated from polysome fractions of exponentially growing Δ*mnmA* cells under untreated and $H_2O_2$-treated conditions and analyzed by mRNA-seq (*n* = 3). (A) Volcano plot showing differentially expressed polysome-associated transcripts in treated Δ*mnmA* cells relative to untreated Δ*mnmA* cells. Significantly upregulated (447) and downregulated (69) genes are indicated. (B) Gene Ontology (GO) enrichment analysis of significantly upregulated transcripts using clusterProfiler. Dot size indicates gene ratio, and color denotes adjusted *P* value. (C) Translational efficiency (TE) plotted against total mRNA changes for treated Δ*mnmA* relative to untreated Δ*mnmA* cells. TE was calculated as the ratio of polysome-bound mRNA to total mRNA, and genes are classified as TE-up (red), TE-down (green), or unchanged (gray). Dashed lines indicate the classification thresholds, and selected genes are labeled. (D) Six-codon signatures of TE-up and TE-down genes in Δ*mnmA* cells following treatment. Average codon Z-scores (±95% CI) are shown for AAA/AAG, GAA/GAG, and CAA/CAG. TE-up genes are shown in red and TE-down genes in green. The dashed vertical line indicates the genome-wide average codon usage after normalization.

**Figure S16**

A

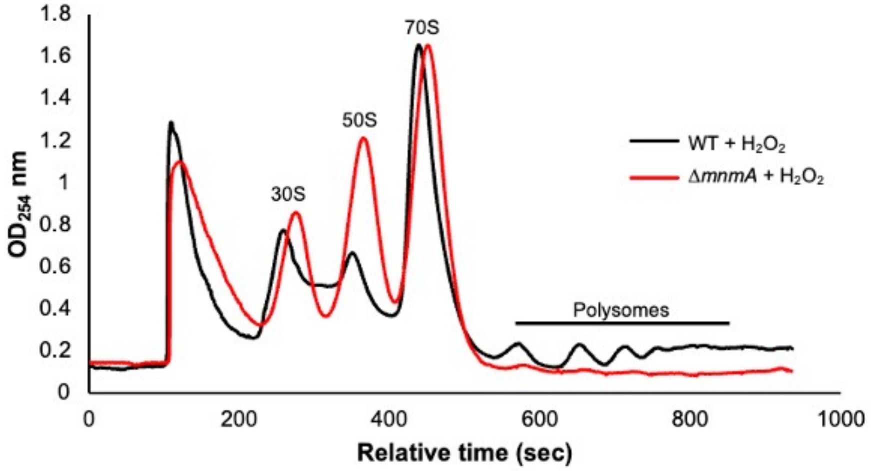


B

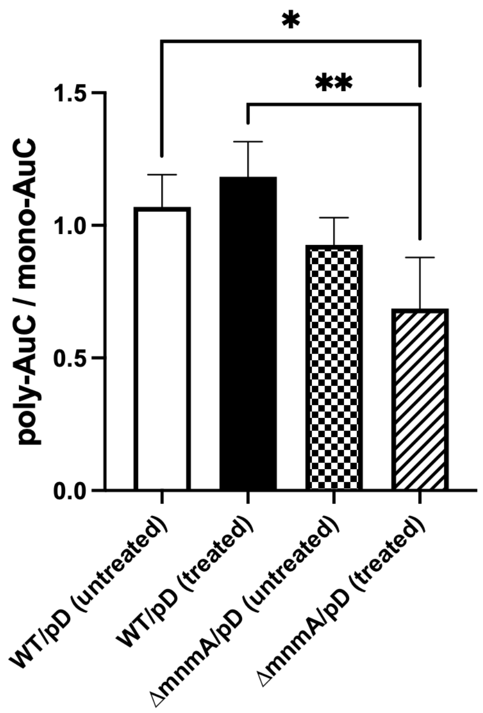


**Supplementary Figure S16. Loss of *mnmA* impairs ribosome engagement and polysome formation under oxidative stress.** (A) Representative sucrose gradient polysome profiles of wild-type (WT) and Δ*mnmA* cells following hydrogen peroxide ($H_2O_2$) treatment. Absorbance at 254 nm ($OD_{254}$) is plotted as a function of fractionation time, with peaks corresponding to the 30S, 50S, and 70S ribosomal species indicated. (B) Quantification of translational activity based on the ratio of polysome-associated ribosomes to monosomes (poly-AUC/mono-AUC) in untreated and $H_2O_2$-treated conditions. $P < 0.05$ (*) and $P < 0.01$ (**). Error bars represent SD from 3 biological replicates.

**Figure S17**

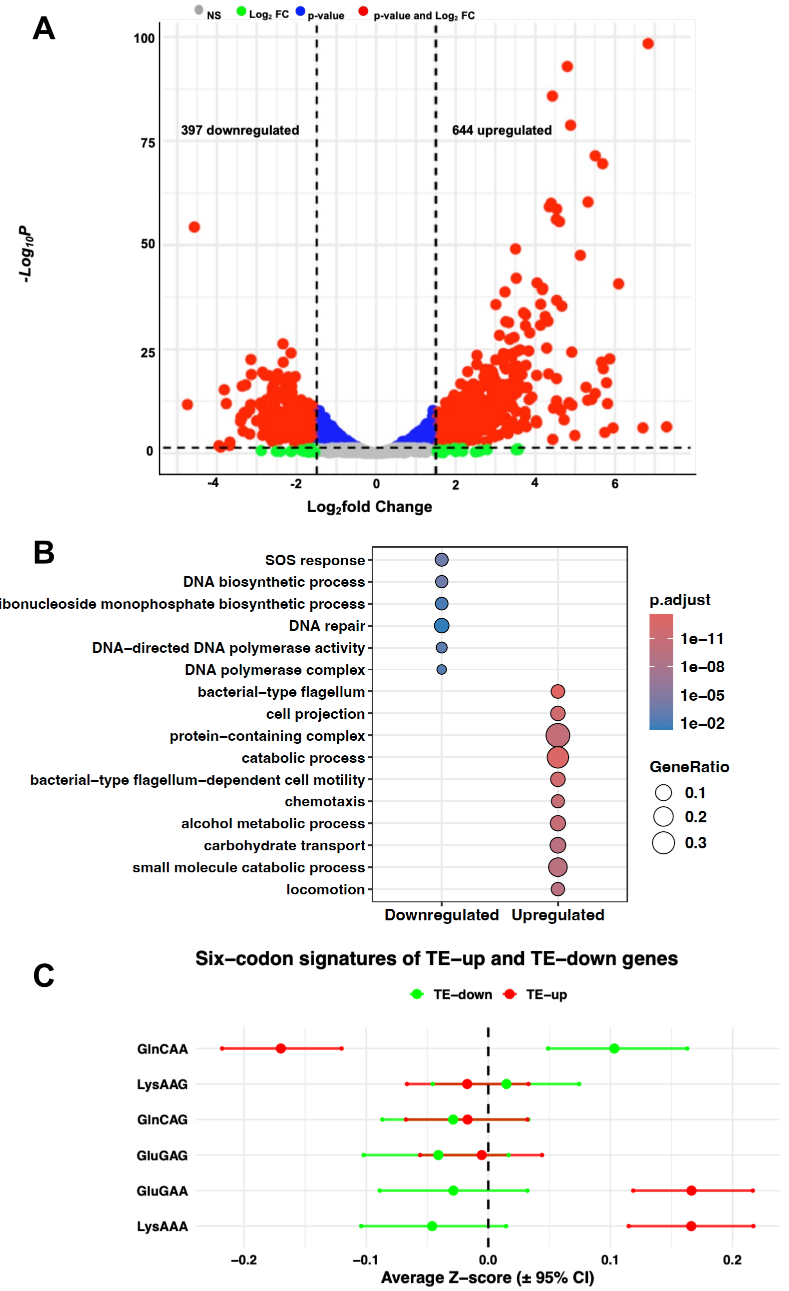


**Supplementary Figure S17. Polysome-associated mRNA profiling of Δ*mnmA* and WT cells under oxidative stress.** Total RNA was isolated from polysome fractions of exponentially growing Δ*mnmA* and WT cells following $H_2O_2$ treatment, and the samples were analyzed by mRNA-seq ($n$ = 3). (A) Volcano plot showing differentially expressed polysome-associated transcripts in treated Δ*mnmA* cells relative to treated WT cells. Significantly upregulated (644) and downregulated (397) genes are indicated. (B) Gene Ontology (GO) enrichment analysis of significantly upregulated and downregulated transcripts using clusterProfiler. Dot size corresponds to the gene ratio, and color indicates the adjusted P-value. (C) Six-codon signatures of TE-up and TE-down genes in treated Δ*mnmA* cells relative to treated WT cells. Average codon Z-scores (±95% CI) are shown for AAA/AAG, GAA/GAG, and CAA/CAG. TE-up genes are shown in red and TE-down genes in green. The dashed vertical line indicates zero, corresponding to the genome-wide average codon usage after normalization.

**Figure S18**

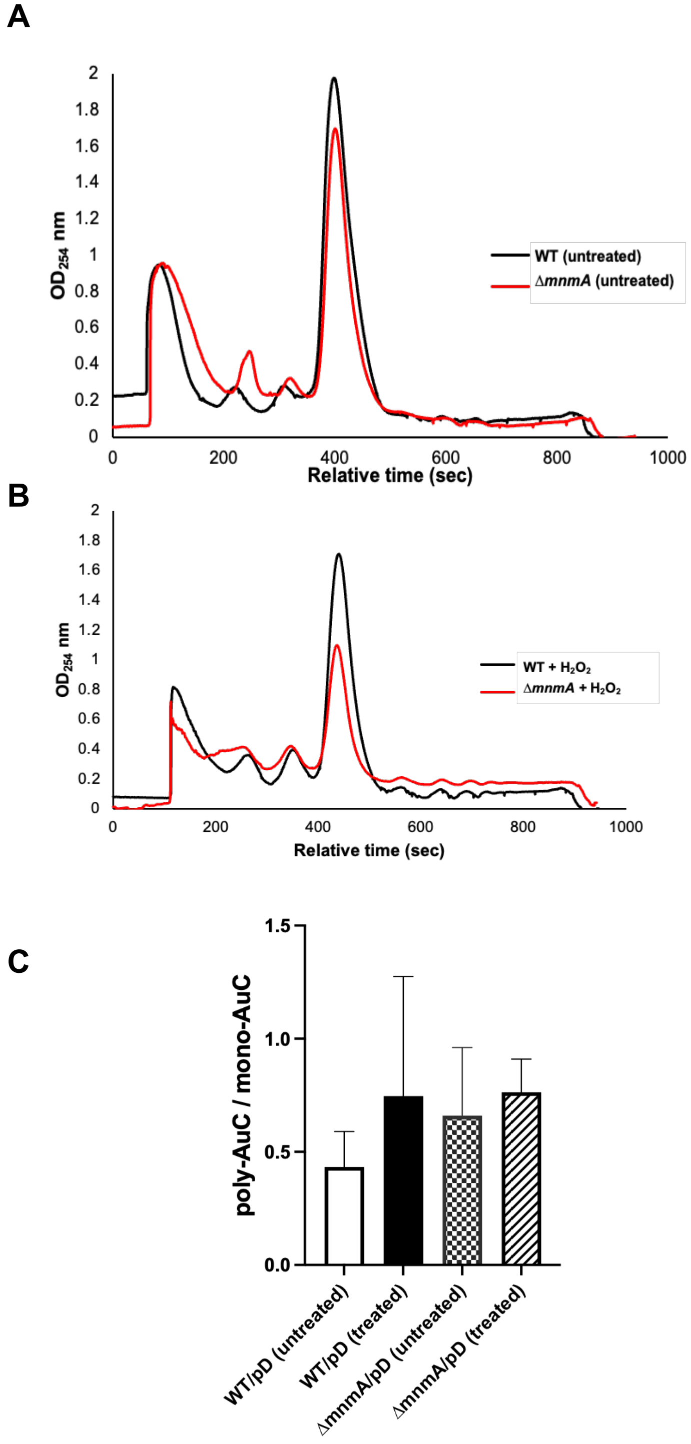


**Supplementary Figure S18: Polysome profile for WT and Δ*mnmA* cells in the stationary growth phase.** Polysome profiling and analysis were performed on WT and ΔmnmA, left untreated or treated with 5 mM $H_2O_2$ for 30 min during the stationary growth phase. (A) Representative polysome traces for WT untreated vs. Δ*mnmA* untreated, and (B) Representative polysome traces for treated WT vs. treated Δ*mnmA*. (C) The polysome-to-monosome area-under-the-curve ratio was analyzed using One-way ANOVA.

Figure S19

A

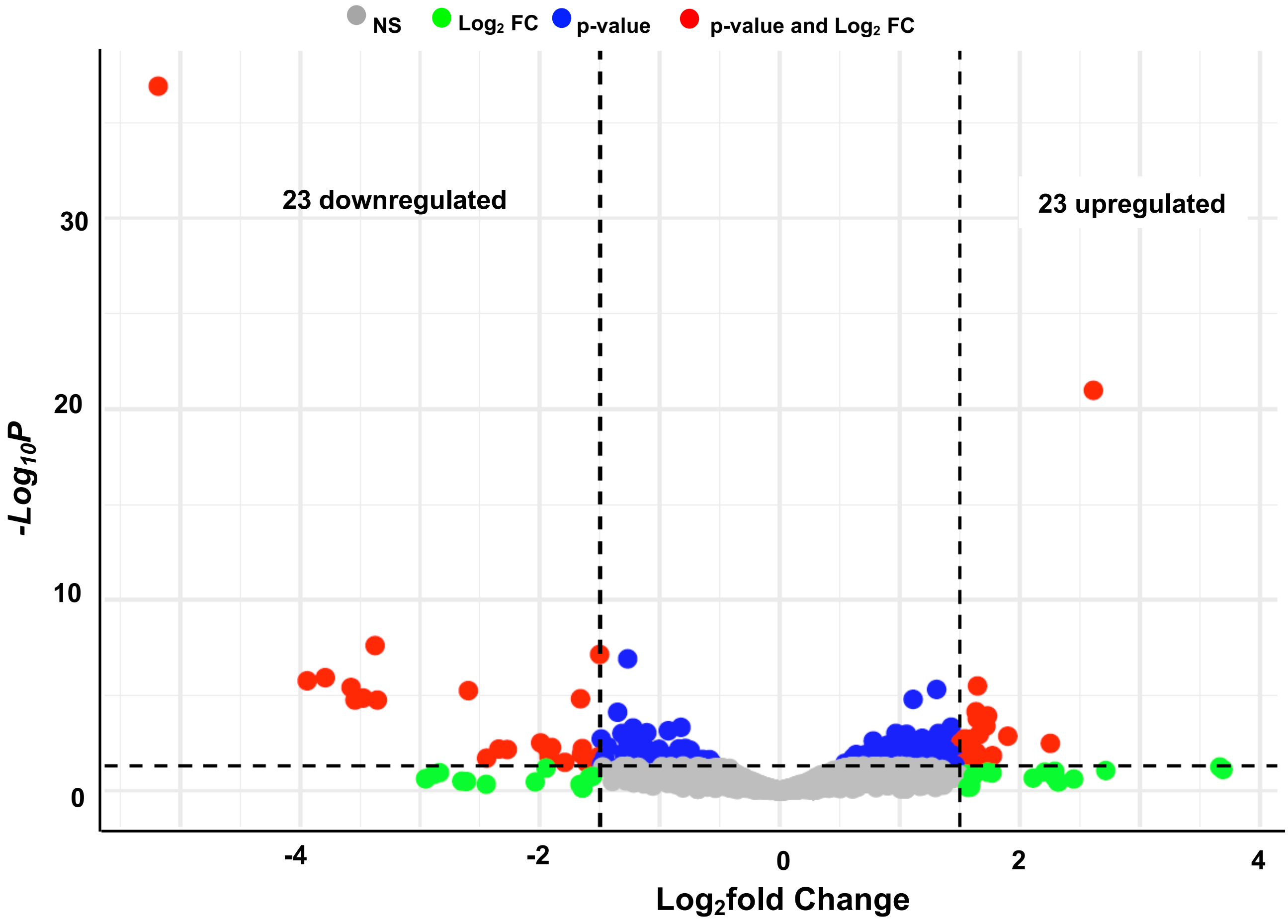


**Supplementary Figure S19: Polysome-associated mRNA analysis for MnmA-deficient and WT cells.** Total RNA from polysome fractions of Δ*mnmA* untreated and WT untreated cells at the stationary growth phase was purified and analyzed by NGS mRNA-seq (n = 3). Volcano plot shows differentially upregulated (23) and downregulated (23) genes.

Figure S20

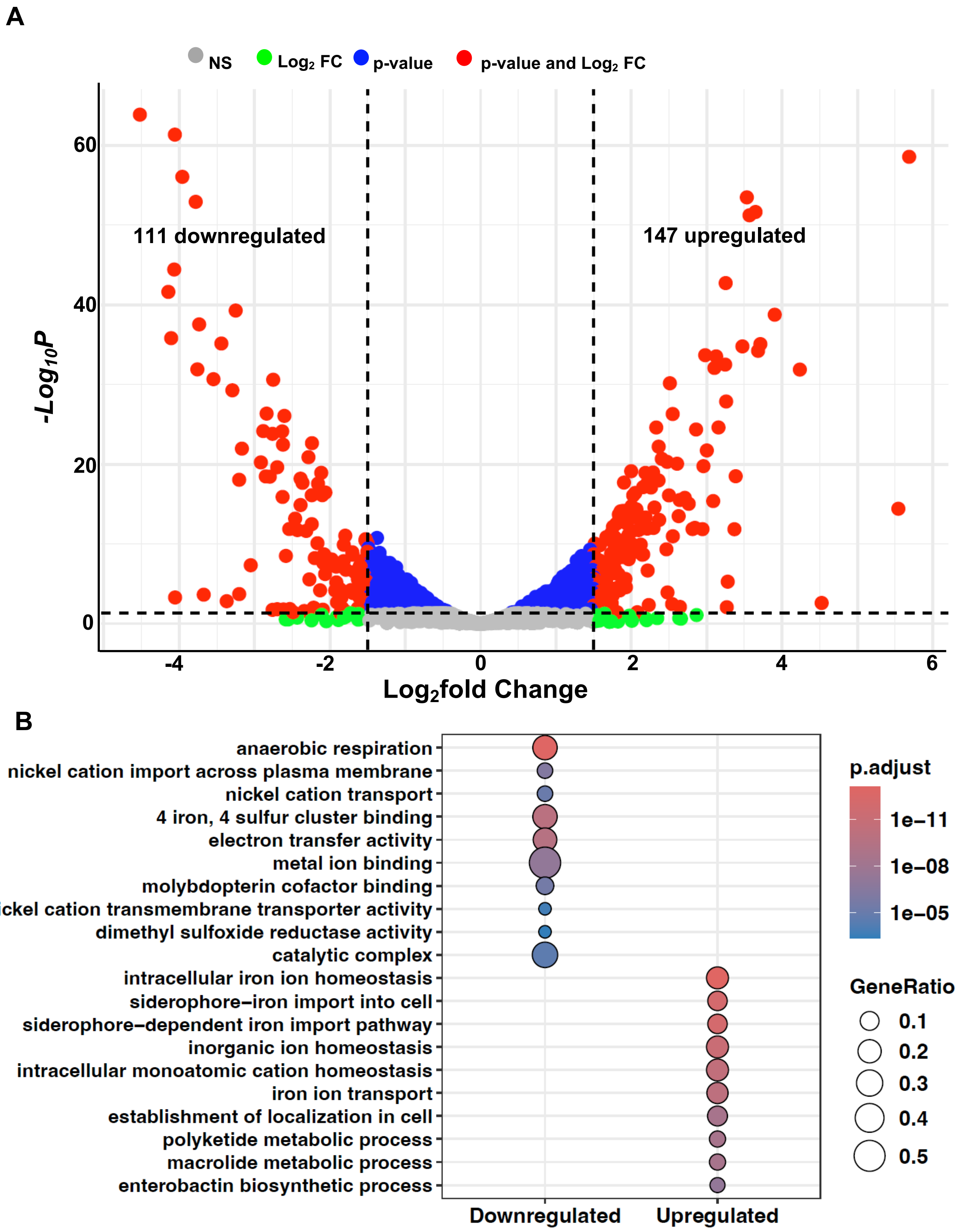


**Supplementary Figure S20: Polysome-associated mRNA analysis for WT cells.** Total RNA from polysome fractions of WT-treated and WT-untreated cells at the stationary growth phase was purified and analyzed by NGS mRNA-seq (n = 3). (A) Volcano plot showing differentially upregulated (147) and downregulated (111) genes. Gene ontology analysis was performed using clusterProfiler for (B) downregulated ($log_2FC \leq 1.50$ and $P_{adj.}$ values $\leq 0.05$) and upregulated genes ($log_2FC \geq 1.50$ and $P_{adj.}$ values $\leq 0.05$).

Figure S21

**A**

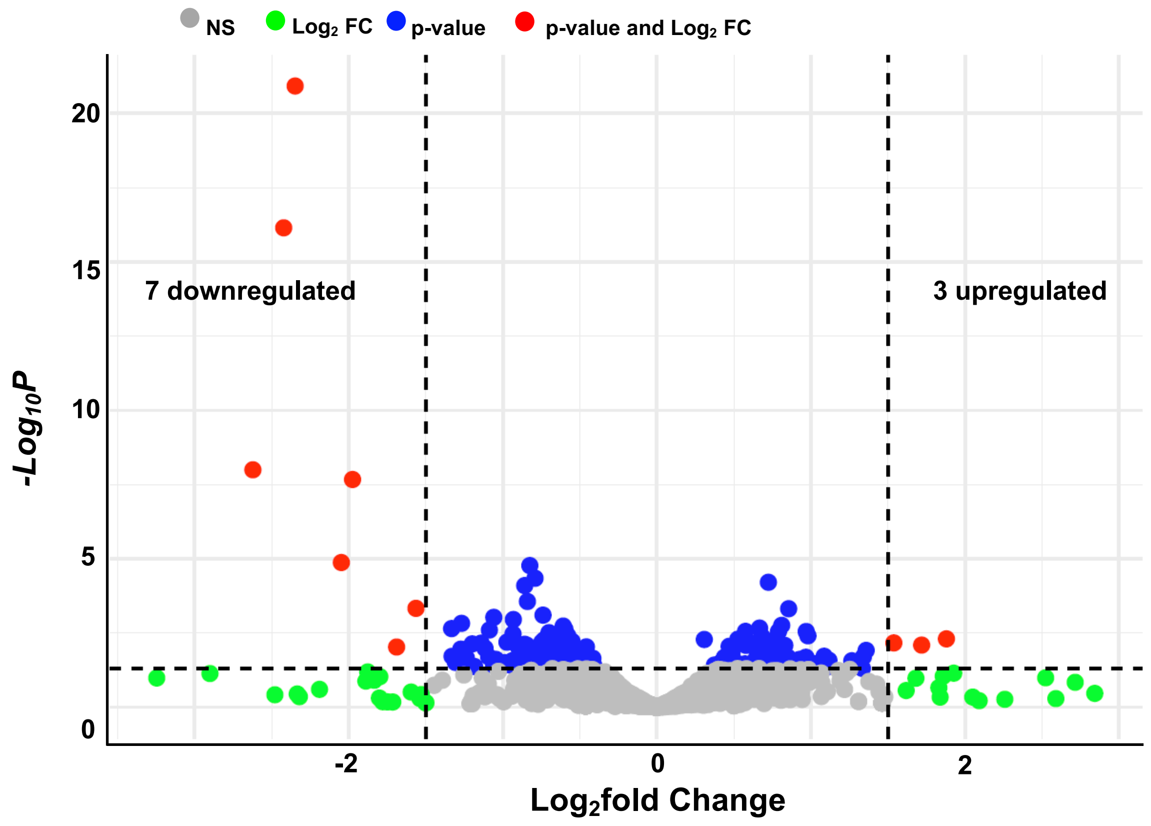


**Supplementary Figure S21: Polysome-associated mRNA analysis for MnmA-deficient cells.** Total RNA from polysome fractions of Δ*mnmA*-treated and Δ*mnmA*-untreated cells at the exponential phase was purified and analyzed by NGS mRNA-seq (n = 3). Volcano plot shows differentially upregulated (3) and downregulated (7) genes.

Figure S22

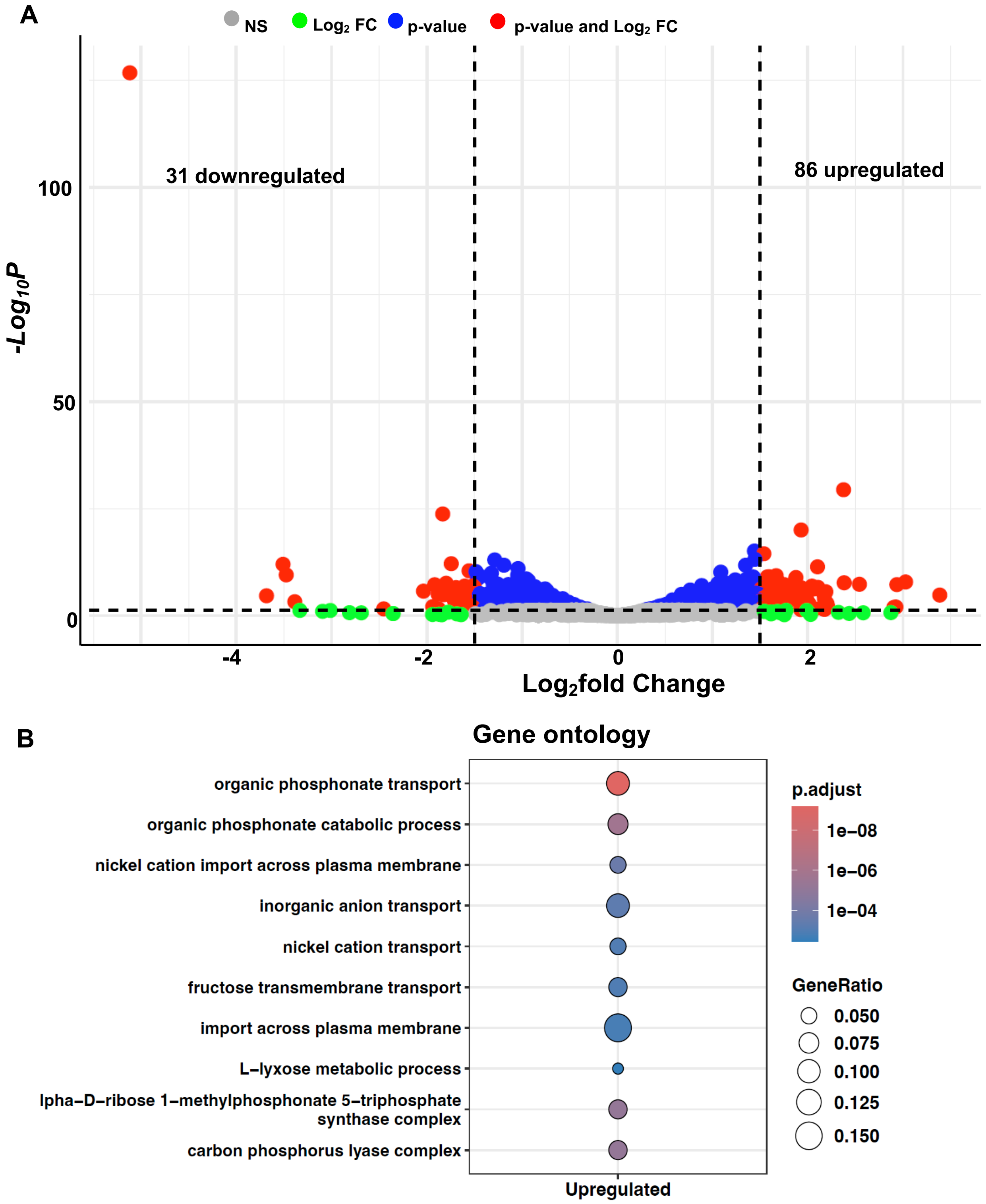


**Supplementary Figure S22: Polysome-associated mRNA analysis for MnmA-deficient and WT cells.** Total RNA from polysome fractions of Δ*mnmA*-treated and WT-treated cells at the exponential phase was purified and analyzed by NGS mRNA-seq (n = 3). (A) Volcano plot showing differentially upregulated (86) and downregulated (31) genes. Gene ontology analysis was performed using clusterProfiler for (B) downregulated ($log_2FC \leq 1.50$ and $P_{adj.}$ values ≤ 0.05) and upregulated genes ($log_2$ FC ≥ 1.50 and $P_{adj.}$ values ≤ 0.05).

**Figure S23**

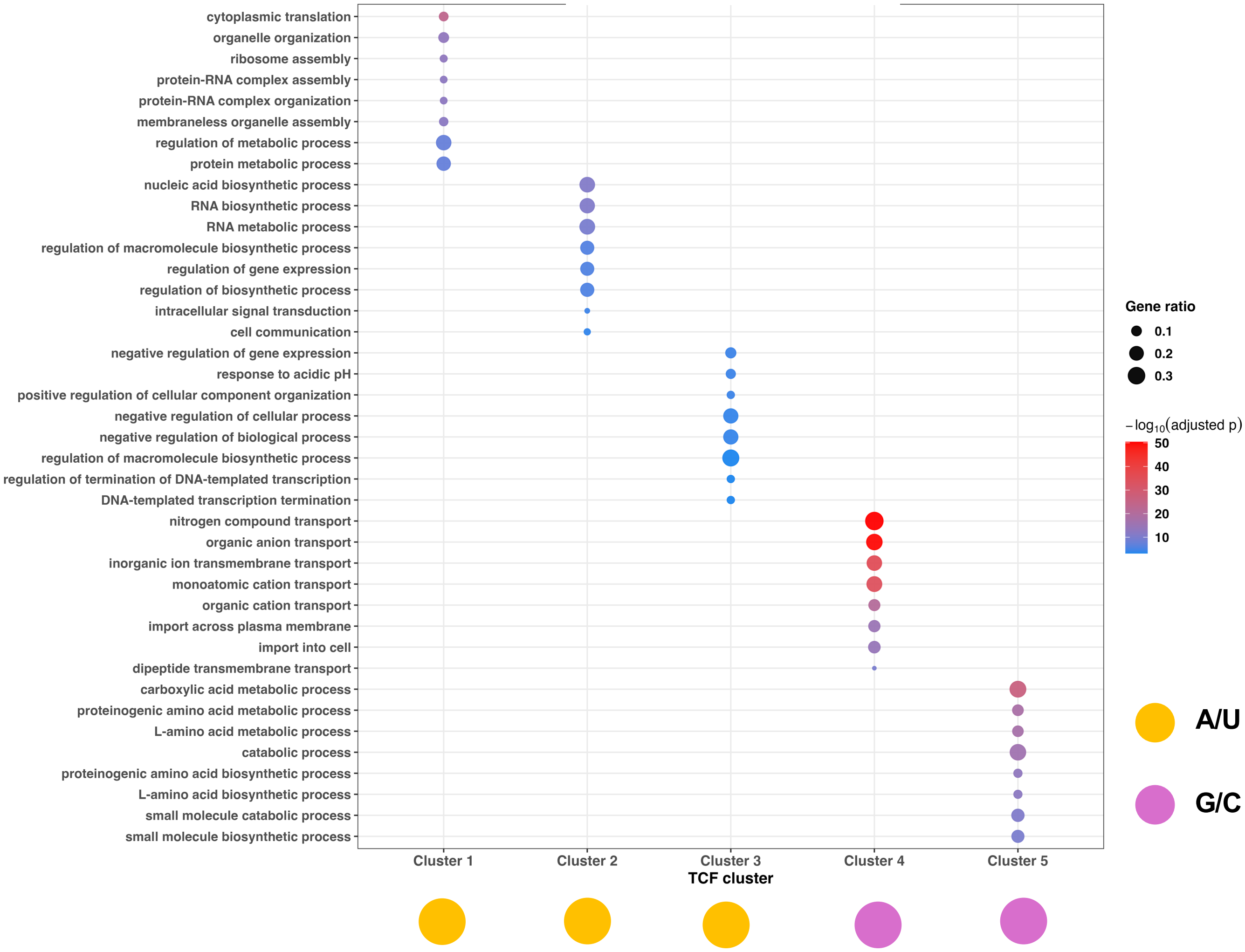


**Supplementary Figure S23. Functional enrichment of codon-defined gene clusters reveals distinct biological programs.** Gene Ontology (GO) biological process enrichment analysis of five gene clusters derived from Gaussian mixture modeling (GMM) of total codon frequency (TCF) Z-score profiles. Each cluster represents a group of genes sharing similar codon usage patterns. Enrichment was performed using clusterProfiler, and results are displayed as a dot plot. Each dot represents a significantly enriched GO term, with dot size corresponding to the gene ratio (the proportion of genes in a cluster associated with the term) and color indicating the adjusted p-value (Benjamini–Hochberg-corrected). Clusters are labeled along the x-axis, and distinct functional signatures are observed across clusters.

**Figure S24**

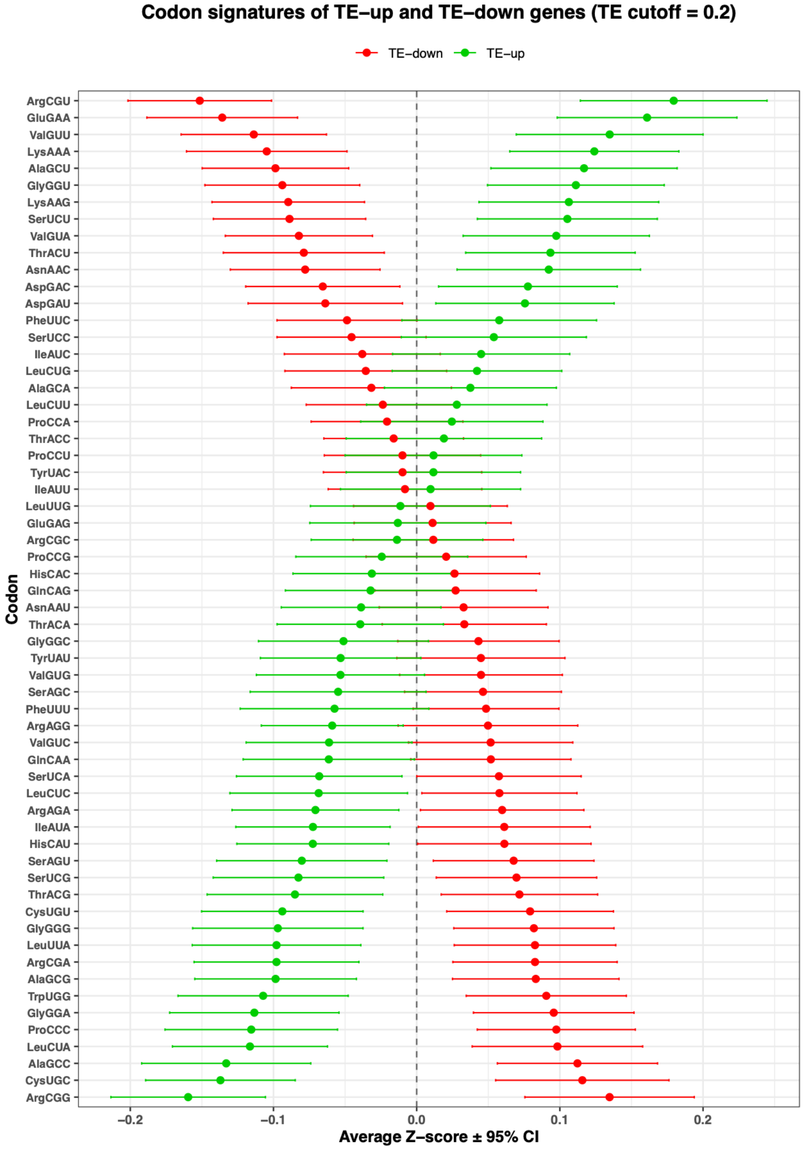


**Supplementary Figure S24. Δ*mnmA* untreated vs WT untreated codon usage signatures associated with translationally upregulated and downregulated genes.** Codon-level enrichment profiles of genes exhibiting increased (TE-up; green) or decreased (TE-down; red) translational efficiency (TE), cutoff = ±0.2. For each codon, frequencies were Z-score normalized across genes, and the mean standardized codon usage ±95% confidence interval was calculated for each translational class. Codons are ordered according to the difference between TE-up and TE-down groups, revealing opposing codon usage patterns between translationally activated and repressed gene sets. Positive values indicate relative enrichment of a codon within a given gene class, whereas negative values indicate relative depletion.

**Figure S25**

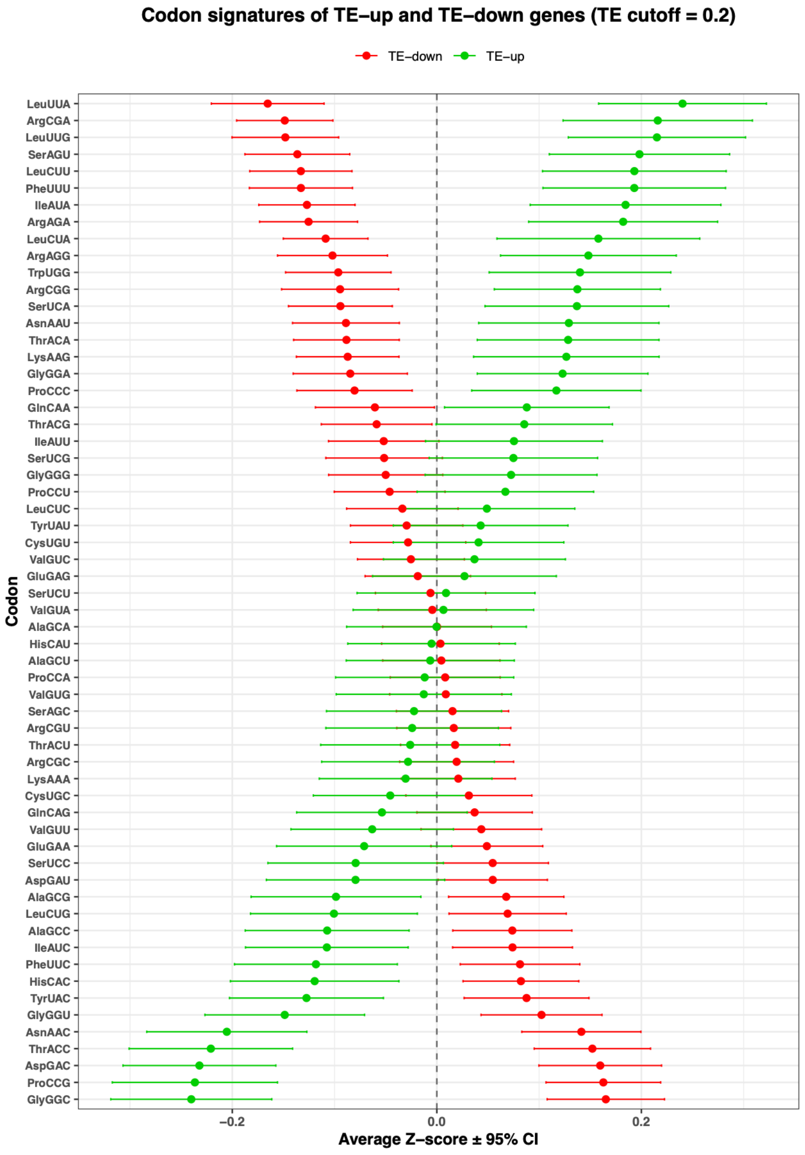


**Supplementary Figure S25. WT-treated vs WT-untreated codon usage signatures associated with translationally upregulated and downregulated genes.** Codon-level enrichment profiles of genes exhibiting increased (TE-up; green) or decreased (TE-down; red) translational efficiency (TE), cutoff = ±0.2. For each codon, frequencies were Z-score normalized across genes, and the mean standardized codon usage ±95% confidence interval was calculated for each translational class. Codons are ordered according to the difference between TE-up and TE-down groups, revealing opposing codon usage patterns between translationally activated and repressed gene sets. Positive values indicate relative enrichment of a codon within a given gene class, whereas negative values indicate relative depletion.

### Figure S26

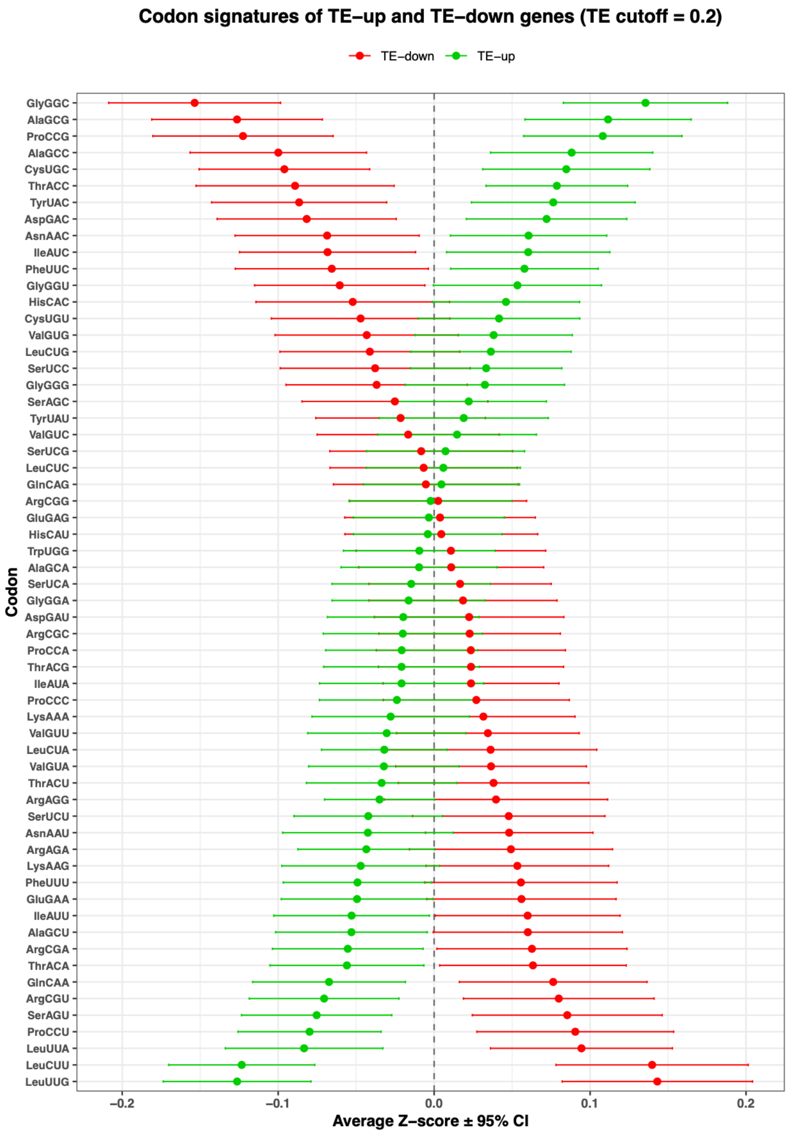


**Supplementary Figure S26. Δ*mnmA*-treated vs Δ*mnmA*-untreated codon usage signatures associated with translationally upregulated and downregulated genes.** Codon-level enrichment profiles of genes exhibiting increased (TE-up; green) or decreased (TE-down; red) translational efficiency (TE), cutoff = ±0.2. For each codon, frequencies were Z-score normalized across genes, and the mean standardized codon usage ±95% confidence interval was calculated for each translational class. Codons are ordered according to the difference between TE-up and TE-down groups, revealing opposing codon usage patterns between translationally activated and repressed gene sets. Positive values indicate relative enrichment of a codon within a given gene class, whereas negative values indicate relative depletion.

### Figure S27

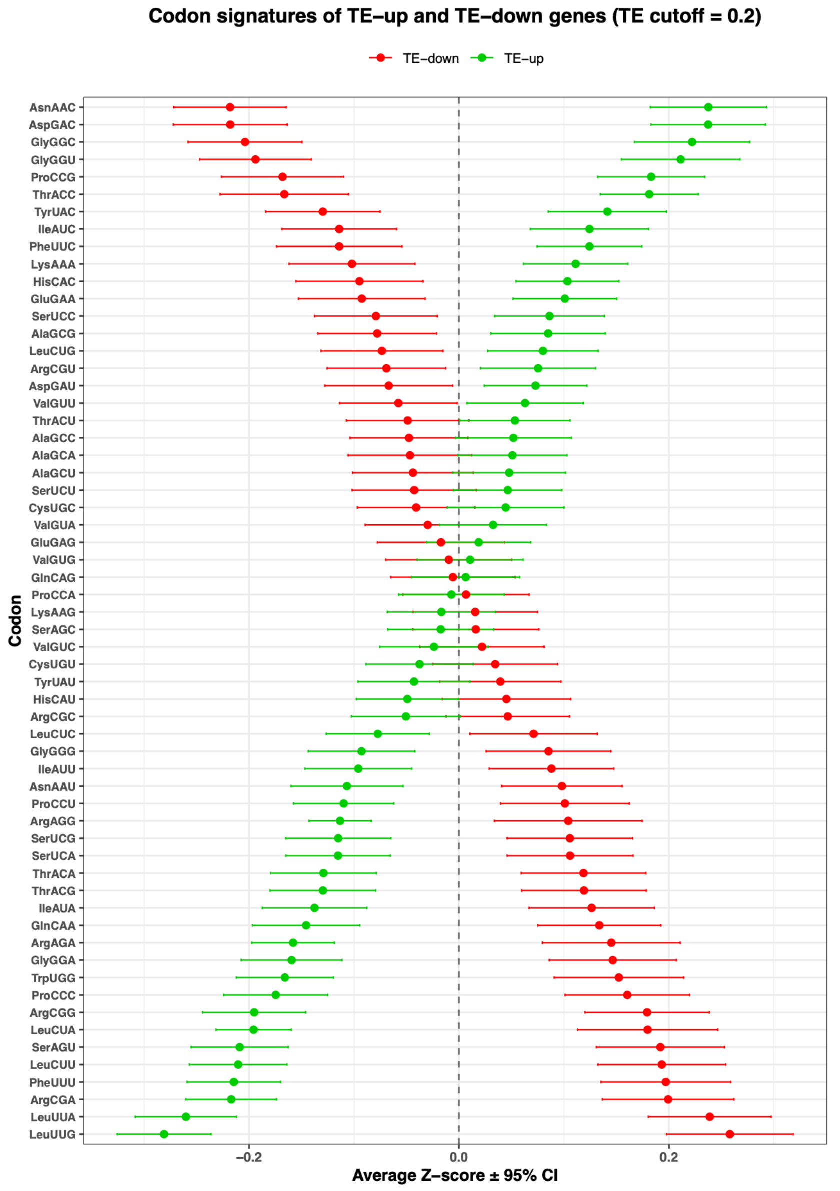


**Supplementary Figure S27. Δ*mnmA*-treated vs WT-treated codon usage signatures associated with translationally upregulated and downregulated genes.** Codon-level enrichment profiles of genes exhibiting increased (TE-up; green) or decreased (TE-down; red) translational efficiency (TE), cutoff = ±0.2. For each codon, frequencies were Z-score normalized across genes, and the mean standardized codon usage ±95% confidence interval was calculated for each translational class. Codons are ordered according to the difference between TE-up and TE-down groups, revealing opposing codon usage patterns between translationally activated and repressed gene sets. Positive values indicate relative enrichment of a codon within a given gene class, whereas negative values indicate relative depletion.

**Table S1**

| GROUPS | Δ*mnmA* untreated vs WT untreated | Δ*mnmA* treated vs Δ*mnmA* untreated | Δ*mnmA* treated vs WT treated | WT treated vs WT untreated |
|---|---|---|---|---|
| **Total regulated genes** | 632 | 385 | 1141 | 913 |
| **Upregulated genes** | 438 | 144 | 575 | 543 |
| **Downregulated genes** | 194 | 241 | 566 | 370 |
| **Total RpoS-regulated genes** | 81 | 34 | 111 | 98 |
| **RpoS-upregulated genes** | 78 | 1 | 18 | 91 |
| **RpoS-downregulated genes** | 3 | 33 | 93 | 7 |
| **Total OxyR-regulated genes** | 8 | 15 | 11 | 18 |
| **OxyR-upregulated genes** | 2 | 14 | 10 | 2 |
| **OxyR-downregulated genes** | 6 | 1 | 1 | 16 |
| **Total LexA-regulated genes** | 0 | 1 | 3 | 1 |
| **LexA-upregulated genes** | 0 | 1 | 2 | 0 |
| **LexA-downregulated genes** | 0 | 0 | 1 | 1 |
| **Total FliA-regulated genes** | 39 | 11 | 43 | 20 |
| **FliA-upregulated genes** | 37 | 1 | 39 | 7 |
| **FliA-downregulated genes** | 2 | 10 | 4 | 13 |
| **Total RpoH-regulated genes** | 18 | 12 | 17 | 21 |
| **RpoH-upregulated genes** | 16 | 3 | 6 | 16 |
| **RpoH-downregulated genes** | 2 | 9 | 11 | 5 |
| **Total RpoN-regulated genes** | 9 | 6 | 12 | 19 |
| **RpoN-upregulated genes** | 6 | 3 | 6 | 13 |
| **RpoN-downregulated genes** | 3 | 3 | 6 | 6 |
| **Total RpoD-regulated genes** | 1 | 7 | 13 | 11 |
| **RpoD-upregulated genes** | 1 | 7 | 13 | 2 |
| **RpoD-downregulated genes** | 0 | 0 | 0 | 9 |
| **Total RpoE-regulated genes** | 0 | 1 | 3 | 1 |
| **RpoE-upregulated genes** | 0 | 0 | 1 | 1 |
| **RpoE-downregulated genes** | 0 | 1 | 2 | 0 |
| **Total FecI-regulated genes** | 0 | 0 | 0 | 0 |
| **FecI-upregulated genes** | 0 | 0 | 0 | 0 |
| **FecI-downregulated genes** | 0 | 0 | 0 | 0 |

**Supplementary Table S1: *E. coli* regulon genes are dysregulated in *E. coli* Δ*mnmA* cells.** Table S1 shows the dysregulated genes for different group comparisons at the exponential phase. The comparisons were made for total and regulon-specific regulated transcripts.

**Table S2**

| GROUPS | ΔmnmA untreated vs WT untreated | ΔmnmA treated vs ΔmnmA untreated | ΔmnmA treated Vs WT treated | WT treated Vs WT untreated |
|---|---|---|---|---|
| **Total regulated genes** | 97 | 372 | 40 | 414 |
| **Upregulated genes** | 35 | 212 | 18 | 214 |
| **Downregulated genes** | 62 | 160 | 22 | 200 |
| **Total RpoS-regulated genes** | 138 | 138 | 138 | 138 |
| **RpoS-upregulated genes** | 5 | 4 | 6 | 5 |
| **RpoS-downregulated genes** | 1 | 16 | 0 | 15 |
| **Total OxyR-regulated genes** | 41 | 41 | 41 | 41 |
| **OxyR-upregulated genes** | 0 | 6 | 0 | 6 |
| **OxyR-downregulated genes** | 3 | 1 | 1 | 1 |
| **Total LexA-regulated genes** | 20 | 20 | 20 | 20 |
| **LexA-upregulated genes** | 0 | 0 | 0 | 0 |
| **LexA-downregulated genes** | 2 | 2 | 0 | 4 |
| **Total FliA-regulated genes** | 74 | 74 | 74 | 74 |
| **FliA-upregulated genes** | 5 | 2 | 3 | 3 |
| **FliA-downregulated genes** | 0 | 6 | 2 | 3 |
| **Total RpoH-regulated genes** | 80 | 80 | 80 | 80 |
| **RpoH-upregulated genes** | 1 | 1 | 0 | 4 |
| **RpoH-downregulated genes** | 0 | 3 | 0 | 7 |
| **Total RpoN-regulated genes** | 73 | 73 | 73 | 73 |
| **RpoN-upregulated genes** | 0 | 10 | 0 | 10 |
| **RpoN-downregulated genes** | 7 | 1 | 4 | 1 |
| **Total RpoD-regulated genes** | 28 | 28 | 28 | 28 |
| **RpoD-upregulated genes** | 0 | 4 | 0 | 4 |
| **RpoD-downregulated genes** | 1 | 2 | 0 | 2 |
| **Total RpoE-regulated genes** | 24 | 24 | 24 | 24 |
| **RpoE-upregulated genes** | 0 | 0 | 0 | 0 |
| **RpoE-downregulated genes** | 0 | 0 | 0 | 0 |
| **Total FecI-regulated genes** | 3 | 3 | 3 | 3 |
| **FecI-upregulated genes** | 0 | 3 | 0 | 2 |
| **FecI-downregulated genes** | 0 | 0 | 0 | 0 |

**Supplementary Table S2: *E. coli* regulon genes are dysregulated in *E. coli* ∆*mnmA* cells.** Table S2 shows the dysregulated genes for different group comparisons at the stationary phase. The comparisons were made for total and regulon-specific regulated transcripts.
